# Molecular inactivation of exopolysaccharide biosynthesis in *Paenibacillus polymyxa* DSM 365 for enhanced 2,3-butanediol production

**DOI:** 10.1101/331843

**Authors:** Christopher Chukwudi Okonkwo, Victor Ujor, Thaddeus Chukwuemeka Ezeji

## Abstract

Formation of Exopolysaccharides (EPS) during 2,3-butanediol (2,3-BD) fermentation by *Paenibacillus polymyxa* decreases 2,3-BD yield, increases medium viscosity and impacts 2,3-BD downstream processing. Therefore, additional purification steps are required to rid the fermentation broth of EPS prior to 2,3-BD purification, which adds to the production cost. To eliminate EPS production during 2,3-BD fermentation, we explored a metabolic engineering strategy to disable the EPS production pathway of *P. polymyxa*, thereby increasing 2,3-BD yield and productivity. The levansucrase gene which encodes levansucrase, the enzyme responsible for EPS biosynthesis in *P. polymyxa*, was successfully disrupted. The resulting *P. polymyxa* levansucrase null mutant showed 34% and 54% increases in growth with 6.4- and 2.4-folds decrease in EPS formation in sucrose and glucose cultures, respectively. The observed decrease in EPS formation by the levansucrase null mutant may account for the 27% and 4% increase in 2,3-BD yield, and 4% and 128% increases in 2,3-BD productivity when grown on sucrose and glucose media, respectively. Genetic stability of the levansucrase null mutant was further evaluated. Interestingly, the levansucrase null mutant remained genetically stable over fifty generations with no observable decrease in growth and 2,3- BD formation with or without antibiotic supplementations. Collectively, our results show that *P. polymyxa* levansucrase null mutant has potential for improving 2,3-BD yield, and ultimately, the economics of large-scale microbial 2,3-BD production.

## Introduction

Considering the finite nature of fossil fuels, recurrent instability in oil price and the environmental concerns associated with oil consumption, there is an urgent need to develop sustainable alternatives to fossil fuels and their derivatives. Over the past few decades, significant attention has been devoted to the development of alternative sources of fuels and chemicals. 2,3- Butanediol (2,3-BD) is an industrial platform chemical that is generated via cracking of petroleum-derived hydrocarbons (e.g. butenes). 2,3-BD has wide industrial applications. For instance, 2,3-BD can be used as a feedstock chemical in the production of 1,3-butadiene (1,3-BD), the monomer from which synthetic rubber is produced (1, 2). Also, 2,3-BD can be used as a feedstock for producing methyl ethyl ketone (MEK), a fuel additive which has a higher heat of combustion than ethanol, and as a solvent from which resins and lacquers can be produced (1, 2). Additionally, 2,3-BD has massive potential as a feedstock for the synthesis of a host of numerous pharmaceuticals, cosmetics, paints, and food preservatives (3, 4).

Several microorganisms have been shown to possess the metabolic machinery to convert carbohydrates to 2,3-BD. However, 2,3-BD is produced via a mixed acid fermentation pathway where other products such as ethanol, acetoin, lactic, formic and acetic acids in addition to exopolysaccharides (EPS) are co-generated. These co-products compete with 2,3-BD for substrates and pyruvate resulting in decreased 2,3-BD yield (5, 6). Several studies have focused on the manipulation of fermentation medium and conditions as means of reducing the accumulation of competing co-products during 2,3-BD fermentation (7, 8, 9, 10, 11). Although, significant progress has been made, accumulation of competing co-products remains a significant challenge to large-scale production of 2,3-BD. This stems from the fact that considerable levels of co-products are still accumulated in the fermentation broth during 2,3-BD fermentation. Further, genetic manipulation of 2,3-BD producers has been explored previously to inactivate lactate dehydrogenase, alcohol dehydrogenase and pyruvate-formate lyase genes - essential genes that encode enzymes involved in the biosynthesis of lactate, ethanol and formic acids, respectively (5, 12, 13, 14). Nevertheless, majority of these studies were conducted with pathogenic 2,3-BD producers which are not ideal for industrial-scale biotechnological applications, as they pose significant health hazards to humans. Thus, we focused on genetic manipulation of *Paenibacillus polymyxa*, a non-pathogenic 2,3-BD producer. *P. polymyxa* was specifically chosen for this study due to its non-pathogenicity and the ability to synthesize levo-2,3-BD, the more desirable 2,3-BD isomer owing to its excellent optical attributes that allow it to be easily dehydrated to 1,3-BD (15, 16). The other 2,3-BD isomers are meso- and dextro-2,3-BD, which are the major fermentation products of the predominantly pathogenic 2,3-BD producers such as *Klebsiella* spp, *Enterobacter aerogenes*, and *Serratia marcescens* (1, 2).

During 2,3-BD fermentation, *P. polymyxa* synthesizes the exopolysaccharide, levan; a fructose polymer with numerous fructose units in β-(2, 6)-linkages (17). Typically, *P. polymyxa* produces more than 50 g/L EPS during fermentation (9), and this accounts for about 20% of the total consumed carbon. Consequently, EPS biosynthesis reduces 2,3-BD titer and yield by diverting carbon away from 2,3-BD biosynthesis. In addition, EPS formation during 2,3-BD fermentation constitutes a major nuisance by clogging of reactor lines which affects proper mixing of fermentation broth, and most importantly, complicates 2,3-BD downstream processing. Additional purification steps would be required to remove EPS prior to 2,3-BD extraction at industrial-scale, which ultimately adds to the overall cost of production. Collectively, reduction in 2,3-BD yield due to EPS formation and the attendant impact on downstream processing adversely affect the economics of 2,3-BD fermentation. Consequently, it is vital to abolish EPS biosynthesis in *P. polymyxa* with a view to re-directing substrate carbons to 2,3-BD biosynthesis for improved titer and yield.

Levan is the only known and characterized EPS synthesized by *P. polymyxa*. Levansucrase plays a key role in levan production in *P. polymyxa* by serving as a conduit for the transfer of fructosyl residues to a growing levan chain (18). *P. polymyxa* produces EPS as a means of attachment to plant roots, the natural habitat of this microorganism (19, 20, 21). Comparative analysis of nucleotide and protein sequences of *P. polymyxa* DSM 365 levansucrase relative to other strains of *P. polymyxa* whose genomes have been completely sequenced and annotated was performed to ascertain the number of copies of levansucrase present in *P. polymyxa* DSM 365. Comparisons were conducted due to absence of complete genome information on *P. polymyxa* DSM 365. The results of this study are shown in **Table 1**. *P. polymyxa* DSM 365 possesses a single copy of levansucrase gene with an open reading frame of 1497 bp. Towards eliminating EPS formation during 2,3-BD fermentation, levansucrase gene was targeted for inactivation in *P. polymyxa*. Using homologous recombination, we report a pioneer work on the inactivation of levansucrase gene of *P. polymyxa*. The *P. polymyxa* levansucrase null mutant developed in this study was evaluated for growth, 2,3-BD production, substrates consumption, 2,3-BD yield and productivity. Further, stability of the levansucrase null mutant was evaluated.

**Table 1.** Comparison of protein and nucleotide sequences between *P. polymyxa* DSM 365 levansucrase and other *P. polymyxa* strains with complete genome sequence using NCBI Blastp and Blastn algorithms for alignment.

| **S/N** | **Levansucrase** | **% Protein identity to *P. polymyxa* DSM 365** | **% Nucleotide sequence identity to *P. polymyxa* DSM 365** | **Accession number** |
| --- | --- | --- | --- | --- |
| 1 | *P. polymyxa* SC2 | 95 | 92 | NC_014622.2 |
| 2 | *P. polymyxa* E681 | 97 | 96 | NC_024483.2 |
| 3 | *P. polymyxa* M1 | 95 | 92 | NC_017542.1 |
| 4 | *P. polymyxa* CR1 | 97 | 96 | NC_023037.2 |

## Materials and methods

### Microorganisms and culture conditions

#### Paenibacillus polymyxa

DSM 365 used in this study was procured from the German Collection of Microorganisms and Cell Culture, Braunschweig, Germany (DSMZ-Deutsche Sammlung von Mikroorganismen und Zellkulturen). The lyophilized stock was reactivated by inoculating into Luria Bertani (LB) broth, grown overnight (12 h), and then stored as glycerol stock (50 % sterile glycerol) at – 80°C. The microorganisms, vectors and enzymes used in this study are shown in **Table 2**.

**Table 2.**
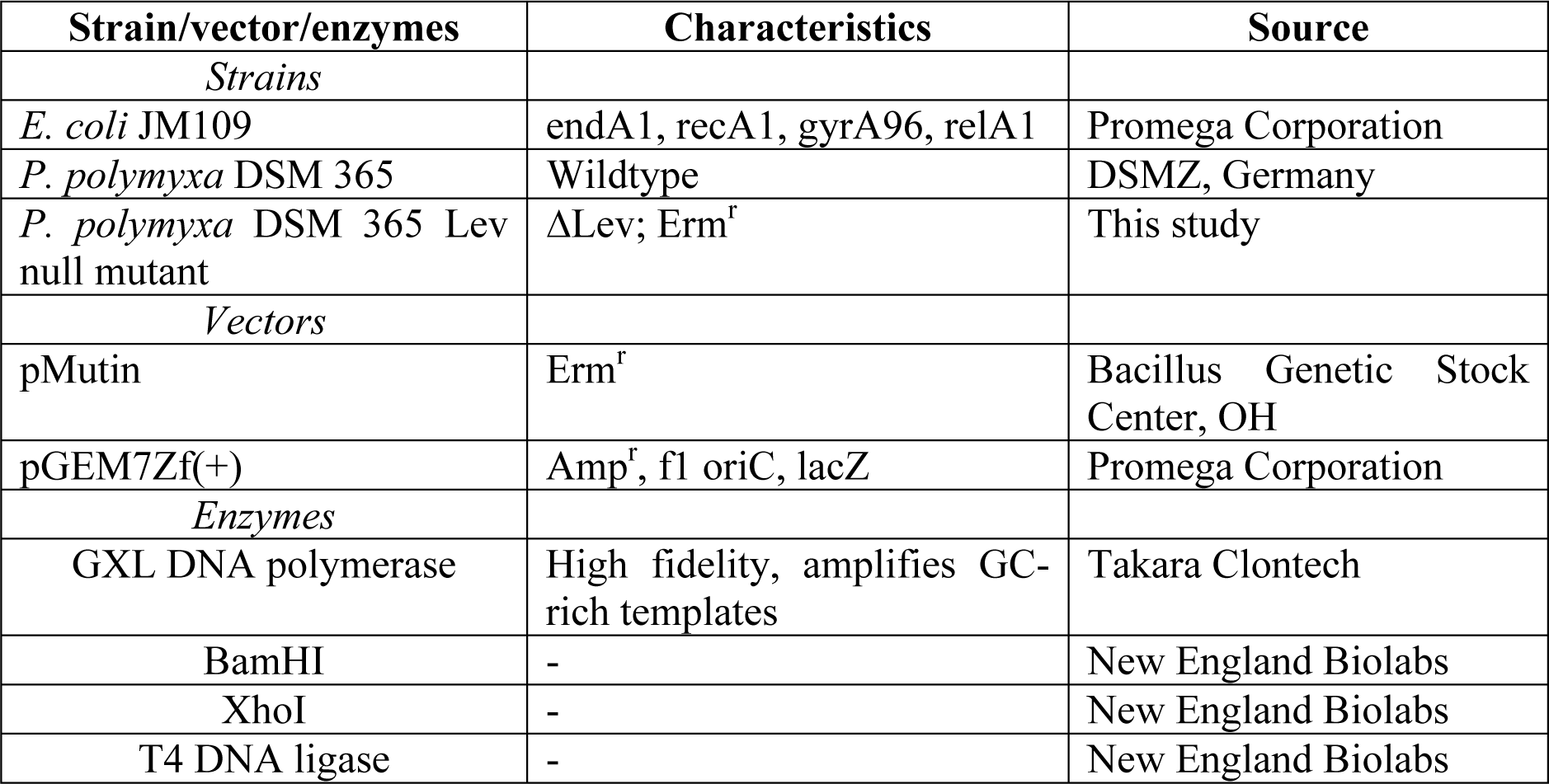
List of microorganisms, vectors and enzymes used in this study and their respective characteristics and sources

### Genomic DNA extraction and amplification of levansucrase inactivation constructs

To extract genomic DNA (gDNA), *P. polymyxa* cells were grown in a previously described pre-culture medium (7) to cell OD_600nm_ of 0.7. The cells were harvested by centrifugation at 10,000 × g and 4°C for 10 min and then, suspended in Tris-HCl-EDTA (TE) buffer (10 mM Tris-HCl, 1 mM EDTA, pH 8.0). Zirconia/Silica beads (0.1 mm, BioSpec Products, Inc., Bartlesville, OK) were added to the cells to a final concentration of 50% (w/v). The cells in the mixture were lysed using a TissueLyzer LT (Qiagen, Hilden, Germany) at 50 oscillations per seconds for 2 min. The cell lysate was centrifuged at 10,000 × g for 10 min and the supernatant was transferred to a clean Eppendorf tube. Phenol-chloroform gDNA extraction method (22) was used to isolate *P. polymyxa* gDNA and then washed with 70% (v/v) ethanol. The gDNA was air-dried at room temperature and re-constituted in 20 μl of nuclease-free water. The gDNA was stored at -20°C until use.

PCR primers for levansucrase gene were designed to amplify the entire levansucrase gene of *P. polymyxa* with the incorporation of *Xho*I and *BamH*I restriction sites at the appropriate locations. The design was such that the PCR primers would amplify short sequences (∼210 bp) upstream and downstream levansucrase gene designated as LevFragA and LevFragB, respectively. Primers used to generate the constructs and their characteristics are shown in **Table 3**. First, the entire levansucrase gene was amplified from the genomic DNA of *P. polymyxa* DSM 365 using LevFragA_fwd and LevFragB_rev primer pair. Then, LevFragA and LevFragB gene fragments were amplified using Lev-FragA_fwd and LevFragA_rev, and LevFragB_fwd and LevFragB_rev, respectively, using gel-purified levansucrase gene amplicon as template. The erythromycin gene was amplified from the plasmid, pMutin (BGSC, Columbus, OH), with primers (Erm_fwd, Erm_rev1 and Erm_rev2; **Table 3**) designed to incorporate ribosomal binding site, spacer and transcription termination sequences. Erm_fwd and Erm_rev1 were first used to amplify erythromycin gene from pMutin. The PCR product was gel-purified and was re-amplified using Erm_fwd and Erm_rev2 primer pair. The use of Erm_rev2 primer in the second amplification of erythromycin gene ensures complete addition of the entire transcription termination sequence downstream of the erythromycin gene sequence. PCR and gene splicing by overlap extension using PCR or gene SOEing (SOEing-PCR) were carried out in a Bio-Rad iCycler™ Thermal Cycler (Bio-Rad, Hercules, CA) using PrimeStar^®^ GXL DNA polymerase (Clontech-Takara, Mountain View, CA). A 50 μl reaction mix containing 5X PrimeStar^®^ GXL buffer (10 μl), dNTPs (0.25 mM), primers (0.5 μM each), DNA template (∼5 ng/ μl) and GXL DNA polymerase (1 μl) was used. The PCR reaction was run using the following conditions: (1) initial denaturation, 98°C for 2 min; (2) 98°C for 20 s (1 cycle); (2) 98°C for 30 s, annealing temperature of primers for 30 s, 72°C for 1 min (35 cycles); (3) final extension, 72°C for 10 min; (4) hold, 4°C for 10 min (1 cycle). Nested PCR was used for one-step SOEing-PCR reaction with the following conditions: (1) initial denaturation, 98°C for 2 min; (2) 98°C for 30 s, annealing temperature of templates overlap region for 30 s; 72°C for 30 s (5 cycles); (3) 98°C for 30 s, annealing temperature of primers, 72°C for 30 s (30 cycles); (4) final extension, 72°C for 5 min; (5) hold, 4°C for 10 min.

**Table 3.**
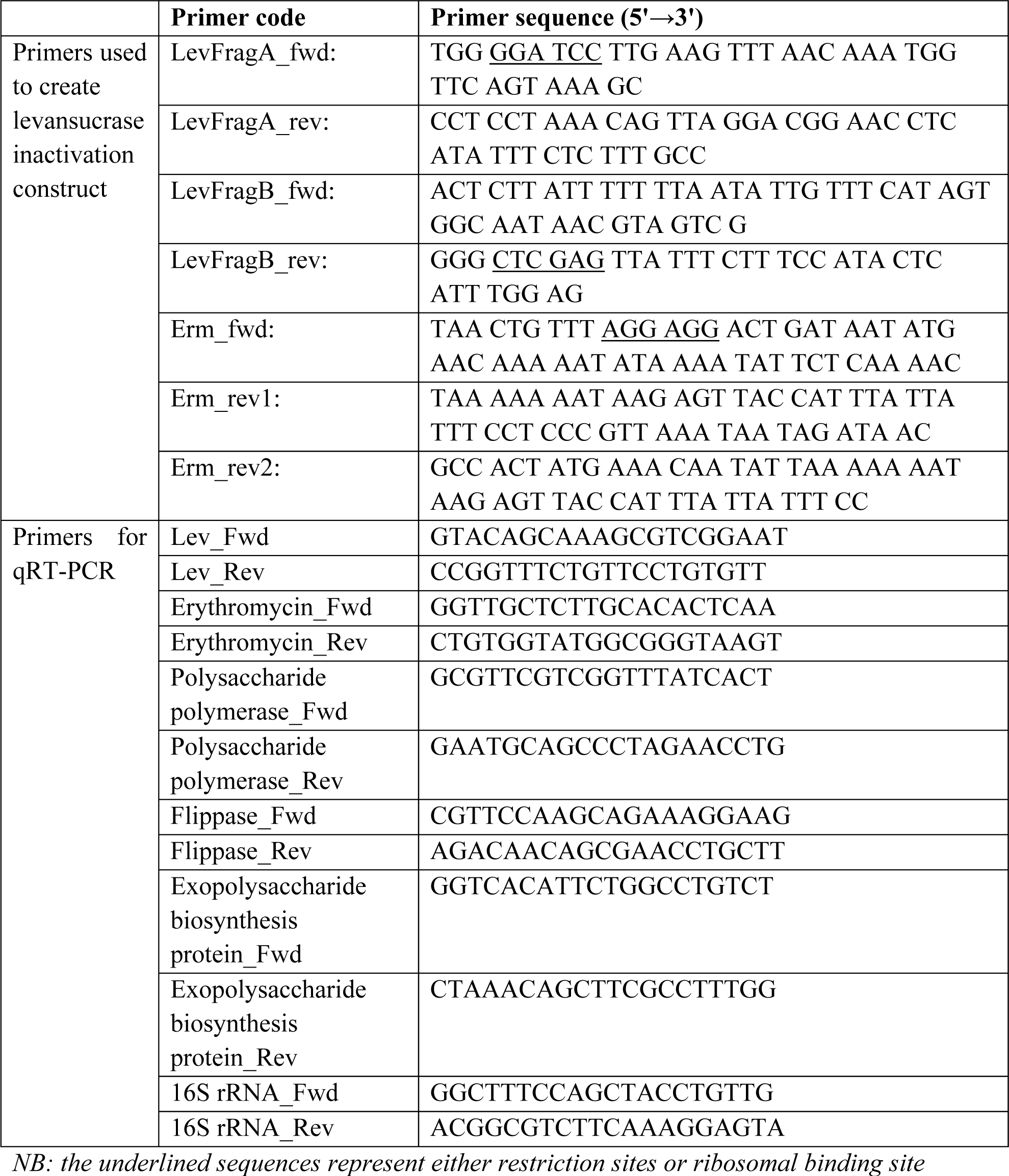
List of primers used to generate levansucrase inactivation construct, and to perform qRT-PCR.

Next, splicing by overlap PCR extension (SOEing-PCR) reactions were used to link LevFragA and ERM genes to generate LevFragA-ERM construct using LevFragA_fwd and Erm_rev2 primer pair and LevFragA and ERM genes as templates. The PCR product (LevFragA-ERM) was gel-purified, and used alongside LevFragB as templates to generate the inactivation construct, LevFragA-ERM-LevFragB, in another SOEing-PCR with LevFragA_fwd and LevFragB_rev primer pair.

### Recombinant plasmid construction

The levansucrase inactivation construct (LevFragA-ERM-LevFragB) was ligated into pGEM^®^7Zf(+), a high copy number plasmid in *E. coli* JM109, which behaves as a non-replicative vector in *P. polymyxa*. pGEM^®^7Zf(+) possesses filamentous phage f1 origin of replication recognized by *E.coli* but not by *P. polymyxa*. Consequently, this vector is used to produce circular single stranded DNA (ssDNA) that enhances homologous recombination *in vivo* (23). The presence of phage f1 origin of replication and the ability of pGEM^®^7Zf(+) to be replicated into stable circular DNA in *E. coli* is important for its application in the inactivation of genes via homologous recombination in *P. polymyxa* and other gram positive bacteria.

pGEM^®^7Zf(+) and LevFragA-ERM-LevFragB were restricted independently with *Xho*I and *BamH*I (New England biolabs, Ipswich, MA) in a 50 μl reaction mixture. The reaction mixture consisted of 5 μl CutSmart buffer (New England biolabs, Ipswich, MA), 1 μl *Xho*I, 0.02 μg/ μl DNA and the reaction volume was made up to 49 μl with nuclease-free water (Amresco^®^, Solon, OH). The mixture was incubated at 37 °C for 1 h, and 1 μl *BamH*I was added and incubated for additional 1 h at 37 °C. The restricted plasmid and LevFragA-ERM-LevFragB construct were purified by agarose gel electrophoresis using GenCatch and advanced PCR extraction kit (Epoch Life Science, Sugar Land, TX). The purified restriction products, LevFragA-ERM-LevFragB and pGEM^®^7Zf(+) were ligated in a 20 μl reaction mix to generate the recombinant pGEM^®^7Zf(+) carrying the levansucrase inactivation construct, LevFragA-ERM-LevFragB. The ligation reaction mixture consisted of 2 μl T4 DNA ligase buffer (New England biolabs, Ipswich, MA), 1 μl T4 DNA ligase (New England biolabs, Ipswich, MA), plasmid (pGEM^®^7Zf(+)) and LevFragA-ERM-LevFragB insert in a ratio of 1:5 with final DNA concentration between 0.02-0.1 pmol. The reaction volume was made up with nuclease-free water (Amresco®, Solon, OH). The reaction mixture was incubated overnight at 16 °C, heat inactivated at 65 °C for 10 min, then chilled on ice for 20 min prior to transformation of competent *E. coli* JM 109 with recombinant pGEM^®^7Zf(+) carrying the levansucrase inactivation construct, LevFragA-ERM-LevFragB. The ligated pGEM^®^7Zf(+) and LevFragA-ERM-LevFragB (recombinant pGEM^®^7Zf(+)) was purified using GenCatch advanced PCR extraction kit (Epoch Life Science, Sugar Land, TX).

### Transformation of competent *E. coli* JM 109

The recombinant pGEM^®^7Zf(+) was used to transform competent *E. coli* JM 109 cells. The recombinant pGEM^®^7Zf(+) (50 ng) was added to 50 μl of competent *E. coli* JM109 cells previously placed on ice for 20 min.. The mixture was heat-shocked at 42 °C for 1 min. Subsequent transformation steps were carried out as previously described (24). The cells were incubated at 37°C and 250 rpm for 1 h after which the cells were plated on LB agar supplemented with 50 μg/ml ampicillin, and 5-bromo-4-chloro-3-indolyl-β-D-galactopyranoside (X-gal) and isopropyl-β-D-1-thiogalactopyranoside (IPTG) to a final concentration of 20 mg/ml and 1 mM, respectively. The plates were incubated at 37°C for 12 h after which white colonies were selected and screened for the presence of correct insert (recombinant pGEM^®^7Zf[+]) by colony PCR and restriction digestion. Colonies with the correct insert were grown in LB medium supplemented with 50 μg/ml ampicillin and the recombinant plasmid was isolated and purified using GenCatch plus plasmid DNA miniprep kit (Epoch Life Science, Sugar Land, TX) and stored at -20 °C prior to use.

### Electro-transformation of competent *P. polymyxa* protoplasts

Following initial unsuccessful attempts to transform competent *P. polymyxa* cells with the recombinant pGEM^®^7Zf(+) harboring the levansucrase inactivation construct via electroporation, competent *P. polymyxa* protoplasts were generated and used instead. Indeed, the cell wall of *P. polymyxa* was removed by a previously described method (25) with slight modifications. Briefly, *P. polymyxa* cells were grown in tryptic soy broth (TSB) for 12 h until cell optical density (OD_600nm_) reached 0.7. The cells were harvested and placed in 50 ml centrifuge tubes pre-chilled on ice for 20 min and then washed twice with 50 mM Tris-Maleate buffer (pH 7.1) containing 2 mM dithiothreitol followed by centrifugation at 1000 × g and 4°C for 7 min. The cell pellets were harvested and re-suspended in Tris-Maleate buffer (pH 7.1) containing 0.6 M sucrose, 5 mM MgCl_2_ and 300 μg/ml lysozyme (Amresco®, Solon, OH). The cell suspension was incubated in an ISOTEMP 220 water bath (Fischer Scientific, Pittsburg, PA) for 60 min at 37°C to make *P. polymyxa* protoplasts. *P. polymyxa* protoplasts were harvested by centrifugation at 1000 × g and 4°C for 7 min. *P. polymyxa* protoplasts were made competent by washing the protoplasts twice with 10% polyethylene glycol (PEG-8000) and re-suspended in 500 μl of 10% PEG-8000. The competent *P. polymyxa* protoplasts were transformed with recombinant pGEM^®^7Zf(+) harboring the levansucrase inactivation construct (LevFragA-ERM-LevFragB) via electroporation. Twenty microliters (100 μg DNA) of the recombinant plasmid was gently mixed with 100 μl of competent protoplasts in a pre-chilled 0.2 cm electroporation cuvette and then placed on ice for 5 min. Electroporation was performed at 2.5 kV, 25 μF capacitance and infinite resistance as previously described (26) in a Bio-Rad Gene Pulser Xcell™ electroporator (Bio-Rad, Hercules, CA). Electric pulse was delivered to the protoplasts between 2.5 and 4.1 milliseconds. Following electroporation, the protoplasts were placed on ice for 5 min and then 500 μl of TSB was added and the mixture was incubated at 35 °C for 6 h to allow protoplast recovery. The recovered cells were plated on tryptic soy agar (TSA) supplemented with 35 μg/ml erythromycin and incubated at 35°C for 16 - 24 h. Colonies were selected and mixed with 50 μl of TSB and then re-plated on a fresh TSA plate supplemented with 35 μg/ml erythromycin and incubated at 35°C for 12 h. Fresh colonies were then selected and colony PCR technique was used to screen for the presence of erythromycin gene. The colonies with erythromycin gene were transferred to TSB supplemented with 50 μg/ml erythromycin. The cells were harvested and genomic DNA was extracted as described above. PCR was performed to screen for the presence of LevFragA-ERM, ERM and ERM-LevFragB fragments using genomic DNA as template. The presence of all three fragments confirmed that levansucrase gene was successfully inactivated via double-cross homologous recombination.

### Characterization of *P. polymyxa* levansucrase null mutant

The *P. polymyxa* levansucrase null mutant was characterized for cell growth, EPS and 2,3-BD production. Batch fermentations were conducted in sucrose- and glucose-based media. One milliliter of 50% glycerol stock of *P. polymyxa* levansucrase null mutant was inoculated into 30 ml of pre-culture medium supplemented with 35 μg/ml erythromycin and incubated at 35 °C and 200 rpm for 6 h until cell optical density (OD_600nm_) reached 1.0-1.2. The actively growing *P. polymyxa* levansucrase null mutant (10%, v/v) was inoculated into the fermentation medium containing 100 g/L sucrose or glucose supplemented with 35 μg/ml erythromycin. The pre-culture and fermentation medium components used in this study have been previously described (7). The fermentation medium was further supplemented with 0 - 0.4 g/L CaCl_2_. Wildtype *P. polymyxa* was prepared as described for the levansucrase null mutant without erythromycin supplementation. Batch 2,3-BD fermentations were conducted in loosely-capped 125 ml Pyrex culture bottles with 30 ml fermentation volume. All experiments were carried out in triplicate and 2 ml samples were collected at 0 h and then, every 12 h until the fermentation terminated. Samples were analyzed for cell growth, culture pH, EPS, 2,3-BD, acetoin, acetic acid and ethanol production.

### Quantitative reverse transcription polymerase chain reaction (qRT-PCR)

To further validate successful deletion of levansucrase gene in *P. polymyxa* genome, the messenger RNA (mRNA) levels of the levansucrase gene was quantified by quantitative reverse transcription PCR (qRT-PCR) using cells grown in sucrose and glucose media. In addition, qRT-PCR was used to confirm successful erythromycin gene integration in *P. polymyxa*. Specific primers for the disrupted region of levansucrase and erythromycin genes were used. Owing to the presence of small amounts of EPS in the *P. polymyxa* levansucrase null mutant cultures, the mRNA levels of a few putative EPS production genes namely flippase, polysaccharide polymerase and exopolysaccharide biosynthesis protein genes were quantified to ascertain their contributions to EPS production on sucrose and glucose substrates. The specific primers used for qRT-PCR are shown in **Table 3**. Total RNA was isolated from the wildtype and levansucrase null mutant using Tri Reagent^®^ (Sigma, St. Louis, MO) following the manufacturer‘s protocol. Culture samples for RNA isolation were taken at 12 h of fermentation (point of maximum EPS accumulation). The RNA content was determined spectrophotometrically using NanoDrop (BioTek^®^ Instrument Inc, Winooski, VT). Total RNA (2 ug) was reverse transcribed to cDNA using Random Hexamers (Qiagen, Hilden, Germany) and M-MLV Reverse Transcriptase (Promega, Madison, WI) according to the manufacturer‘s protocol. The qRT-PCR was conducted in triplicates using the resulting cDNA and GoTaq^®^ qPCR Master Mix containing BRYT Green^®^ (Promega, Madison, WI) in a Bio-Rad CFX96 Touch Deep Well™ Real-Time Detection Systems (Bio-Rad, Hercules, CA). The conditions for the qRT-PCR were: 1 cycle of 95°C at 15 min (initial denaturation), then 40 cycles of 55°C at 30 s (annealing and extension), followed by melting curve analysis via heating from 55°C to 95°C with 1°C per 10 s temperature increment. The mRNA expression levels of all the tested genes were normalized to *P. polymyxa* 16S rRNA (internal standard) and relative expression was performed by the 2^-δδCT^ method (27).

### Analytical methods

Microbial cell growth was determined by measuring its optical density (OD_600_) in a DU^®^ Spectrophotometer (Beckman Coulter Inc., Brea, CA). Changes in pH were measured using an Acumen^®^ Basic pH meter (Fischer Scientific, Pittsburgh, PA). Concentrations of fermentation products, 2,3-BD, acetoin, ethanol, and acetic acid were quantified using a 7890A Agilent gas chromatograph (Agilent Technologies Inc., Wilmington, DE, USA) equipped with a flame ionization detector (FID) and a J × W 19091 N-213 capillary column [30 m (length) × 320 μm (internal diameter) × 0.5 μm (HP-Innowax film thickness)] as previously described (28).

The concentration of sugars such as sucrose, glucose and fructose was quantified by high performance liquid chromatography (HPLC) using a Waters 2796 Bioseparations Module equipped with an Evaporative Light Scattering Detector (ELSD; Waters, Milford, MA) and a 9 μm Aminex HPX-87P column; 300 mm (length) × 7.8 mm (internal diameter) connected in series to a 4.6 mm (internal diameter) × 3 cm (length) Aminex deashing guard column (Bio-Rad, Hercules, CA). The column temperature was maintained at 65°C. The mobile phase was HPLC-grade water (Waters Corporation, Milford, MA) maintained at a flow rate of 0.6 ml/min as described previously (29).

The EPS produced during fermentation was quantified using a method described by (30) with modifications. Culture broth was centrifuged at 8,000 × g for 10 min to pellet the cells while EPS was retained in the supernatant. The EPS was then precipitated with 95 % ethanol (4°C); 10 × the volume of the supernatant. The supernatant-ethanol mixture was kept overnight at 4 °C followed by centrifugation at 8,000 × g for 10 min. The EPS pellet was dried in the oven at 60°C and reconstituted in distilled water. The EPS containing solution was vortexed vigorously to ensure complete dissolution of the EPS. The EPS was then quantified by Phenol-sulfuric acid method (31, 32). Briefly, 25 μl of 80% phenol was added into test tubes A containing 1 ml glucose standards (0.1g/L) and test tubes B containing 1 ml diluted EPS samples. The mixture was vortexed briefly and 2.5 ml concentrated sulfuric acid (Fischer Scientific, Pittsburg, PA) was added to the mixture. The mixture was left to stand for 10 min. The text tubes containing the mixture were incubated at 25°C for 10 min. After incubation, the mixture was gently vortexed and absorbance was measured at 490 nm against reagent blank prepared as the samples. A standard curve was generated by plotting the values of glucose concentration (X-axis) against absorbance (OD_490nm_) (Y-axis) and EPS concentration was interpolated from the standard curve.

### Levansucrase assay

Levansucrase activity in the levansucrase null mutant and wildtype *P. polymyxa* were measured using culture supernatant. Levansucrase activity was determined using a method modified from (33). *P. polymyxa* samples (levansucrase null mutant and wildtype) were collected at the exponential growth phase when maximum EPS is produced. The sample was centrifuged for 20 min at 8,600 × g and 4°C. The supernatant from each sample was divided into two portions. One portion was used to quantify EPS as described above and the EPS obtained was designated as [EPS]_B._ The other portion was used to assay levansucrase activity and the total EPS produced after levansucrase activity assay was designated as [EPS]_A_. The reaction mixture for the levansucrase activity assay consisted of 400 μl of 1 M sucrose in 50 mM phosphate buffer (pH 6.0) and 100 μl of culture supernatant. The mixture was incubated at 35 °C for 1 h. Following levansucrase activity, EPS was precipitated with 95% ethanol (4°C) and the EPS was subsequently quantified and expressed as [EPS]_A_. The EPS produced during levansucrase activity was determined from the equation below:

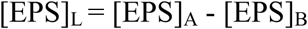

Where [EPS]_L_ represents the concentration of EPS synthesized during levansucrase assay.

The concentrations of protein in the supernatants were determined by Bradford method (34). One unit of levansucrase activity was defined as the milligram of protein that catalyzed the formation of one μmoles of EPS (levan) per min at 35°C in 1 M sucrose solution.

### Growth rate and generation time of levansucrase null mutant

To determine the stability of the levansucrase null mutant, the growth curve was first obtained. Cells were grown to exponential phase (10 h) in the pre-culture medium (7), then harvested and washed twice with sterile distilled water by centrifugation at 5,000 × g for 3 min. The cell pellet was reconstituted to several dilutions and the optical densities (OD_600nm_) were measured against sterile distilled water as blank. Each cell dilution was then centrifuged in a pre-weighed Eppendorf tube at 10,000 × g for 10 min and the supernatant was discarded. The cell pellets were dried for 18 h in TempCon™ Oven (American Scientific Products, McGraw Park, IL) at 50 °C. The cells were weighed and the weight of the cells at each OD_600nm_ reading was determined. A standard curve was generated from a plot of cell biomass (mg/L) against absorbance at OD_600nm_. The standard curve was then used to convert optical density measurements at OD_600nm_ to cell biomass concentrations. The growth curve was obtained by growing the levansucrase null mutant in pre-culture medium and the cell biomass was measured at several time points until the cells reached the death phase of growth. Then, the cell biomass was plotted against time. The generation (doubling) time of levansucrase mutant was determined from the exponential phase of the growth curve (**Fig. S1**).

### Stability of *P. polymyxa* levansucrase null mutant

The stability of levansucrase null mutant was determined in the presence and absence of antibiotic, erythromycin, for 50 generations. The levansucrase null mutant was grown in pre-culture medium supplemented with 35 μg/ml erythromycin until OD_600nm_ reached 1.0-1.2. The actively growing cells (10%, v/v) were transferred into fermentation medium containing 100 g/L sucrose supplemented with 35 μg/ml erythromycin and this generation was regarded as G_0_ (generation zero). The generation time of *P. polymyxa* levansucrase null mutant was pre-determined to be 1.5 h (**Fig. S1**). Several subcultures (every 3 h i.e., 2 generations) were made from G_0_ until G_50_ (generation 50) was attained, and in each case, cultures were supplemented with 35 μg/ml erythromycin. Fermentations were conducted using G_0_, G_10_, G_20_, G_30_, G_40_, and G_50_ under antibiotic selective pressure. Samples (2 ml) were drawn at 0 h and every 12 h until the fermentation ended and then, analyzed for cell growth, EPS and 2,3-BD production. The same experiment was conducted without antibiotic supplementation. Generations G_0_, G_10_, G_20_, G_30_, G_40_, and G_50_ were obtained as described above and then used to conduct fermentations. The antibiotic resistance of all the generations studied (with and without erythromycin supplementation) was determined by PCR and replica plating onto erythromycin-containing plates. For each generation, samples of *P. polymyxa* levansucrase null mutant were drawn during the stationary growth phase (12-16 h of growth) and sub-cultured into fresh pre-culture medium and grown until another stationary growth phase (12-16 h of growth) was attained. The cells were diluted to a concentration of 10^8^ cfu/ml and were plated on TSA plates without antibiotic (erythromycin) supplementation. The plates were incubated at 35 °C for 12 h. Colonies from each generation were selected and screened for presence of the erythromycin gene using PCR. Recombinant pGEM^®^7Zf(+) harboring the levansucrase inactivation construct was used as erythromycin gene control. Gel electrophoresis was performed using the colony PCR products. The presence of erythromycin gene in the colonies confirmed the stability of *P. polymyxa* levansucrase null mutant. Colonies from the TSA plates (without erythromycin supplementation) were transferred by replica plating to fresh TSA plates supplemented with erythromycin (35 μg/ml). The antibiotic-supplemented plates were incubated at 35 °C for 12 h and the resulting colonies were quantitatively compared to the plates without antibiotic supplementation. A schematic representation of step-by-step procedure employed to evaluate the stability of the *P. polymyxa* levansucrase null mutant is shown in **Fig S2**.

### Statistical analysis

General Linear Model of Minitab 17 (Minitab Inc., State College, PA) was used for all statistical analyses. Analysis of variance (ANOVA) using Tukey‘s method for pairwise comparisons was employed to compare differences between treatments. Differences in growth, sugar utilization, maximum product concentrations, 2,3-BD yields and productivities were compared at 95 % confidence interval. 2,3-BD yield was expressed as the gram of 2,3-BD produced from one gram of substrate (sucrose or glucose). 2,3-BD productivity was expressed as the gram per liter of 2,3-BD produced per hour of fermentation.

## Results

### Levansucrase gene inactivation in *P. polymyxa* DSM 365

The levansucrase gene of *P. polymyxa* was successfully inactivated by double-cross homologous recombination. Erythromycin resistance gene was inserted between a 210 bp upstream fragment and a 213 bp downstream fragment of levansucrase gene creating a 1224 bp levansucrase inactivation construct (**Fig 1**). The inactivation construct design included a stop codon downstream of LevFragA sequence followed with a ribosomal binding site and a spacer sequence upstream of erythromycin gene. In addition, the construct included a transcription terminator sequence downstream of the erythromycin gene that precedes the LevFragB sequence. The strategy and design employed in generating levansucrase inactivation construct is shown in **Fig 1**. Inclusion of stop codon, ribosomal binding site and spacer sequence, and transcription terminator sequences ensured that only erythromycin gene is transcribed into mRNA without creating additional metabolic burden on *P. polymyxa*. Following transformation of *P. polymyxa,* PCR-screening of the levansucrase null mutant colonies using LevFragA_fwd/Erm_rev2, Erm_fwd/Erm_rev2 and Erm_fwd/LevFragB_rev primer pairs showed that the mutant possesses LevFragA-ERM, ERM, and ERM-LevFragB genes corresponding to 1008, 800, and 1015 bp, respectively (**Fig 2**), thus confirming successful double-cross homologous recombination.

**Figure 1.**
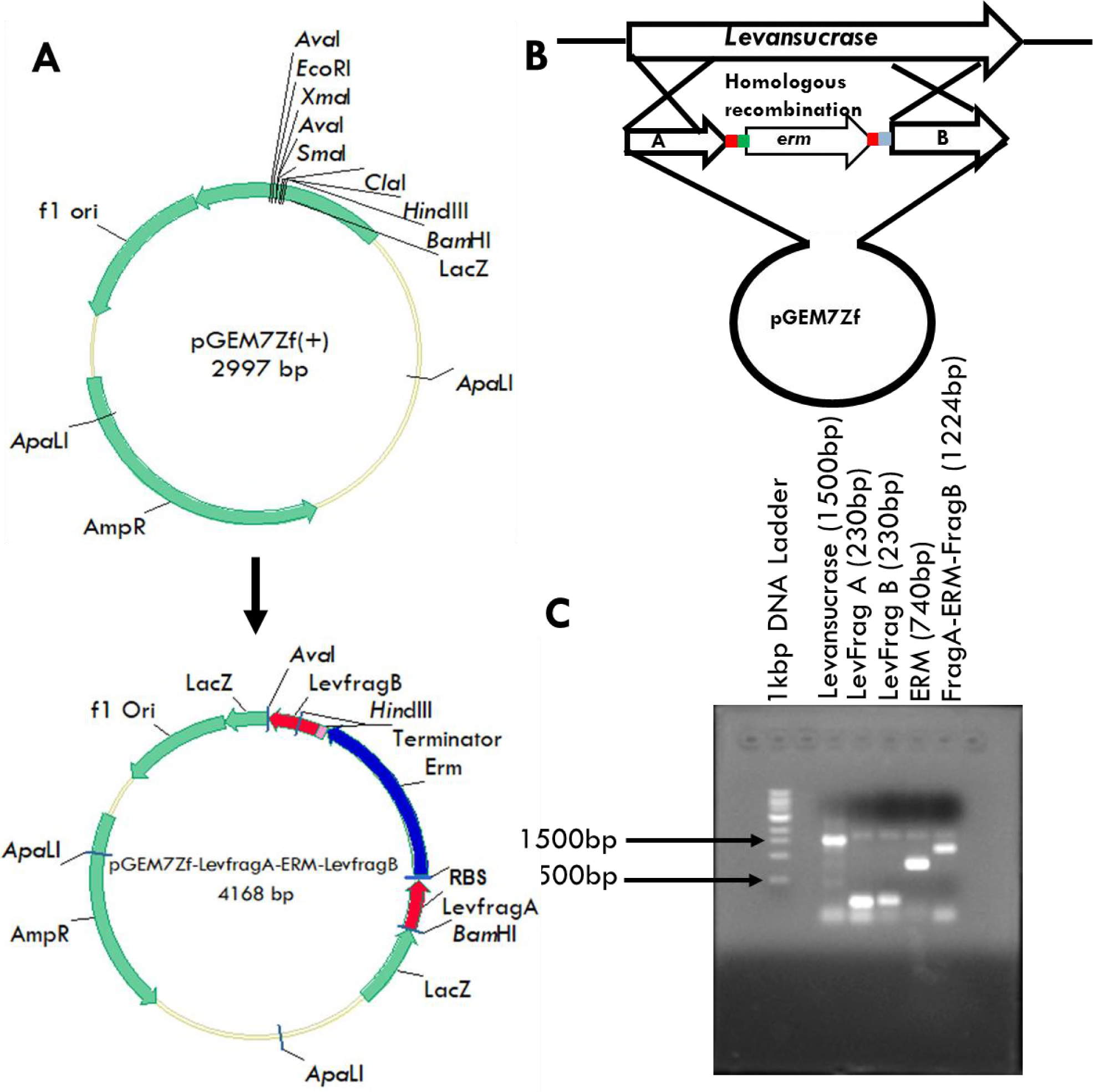
Levansucrase inactivation construct generation. **A**: Recombinant pGEM7Zf-LevfragA-Erm-LevfragB generated from the parent plasmid, pGEM7Zf(+). **B**: levansucrase gene was amplified from the genome of *P. polymyxa* and was used to generate the inactivation construct with erythromycin gene placed between the upstream (210 bp) sequence (LevFragA) and downstream (213 bp) sequence (LevFragB) of levansucrase gene. The construct was ligated into previously double digested pGEM7Zf(+) and used to inactivate levansucrase gene in the chromosome of *P. polymyxa* via double-cross homologous recombination. **C**. Gel image showing levansucrase gene, LevFragA, LevFragB, ERM, LevFragA-ERM and LevFragA-ERM-LevFragB gene fragments during generation of levansucrase inactivation construct.

**Figure 2.**
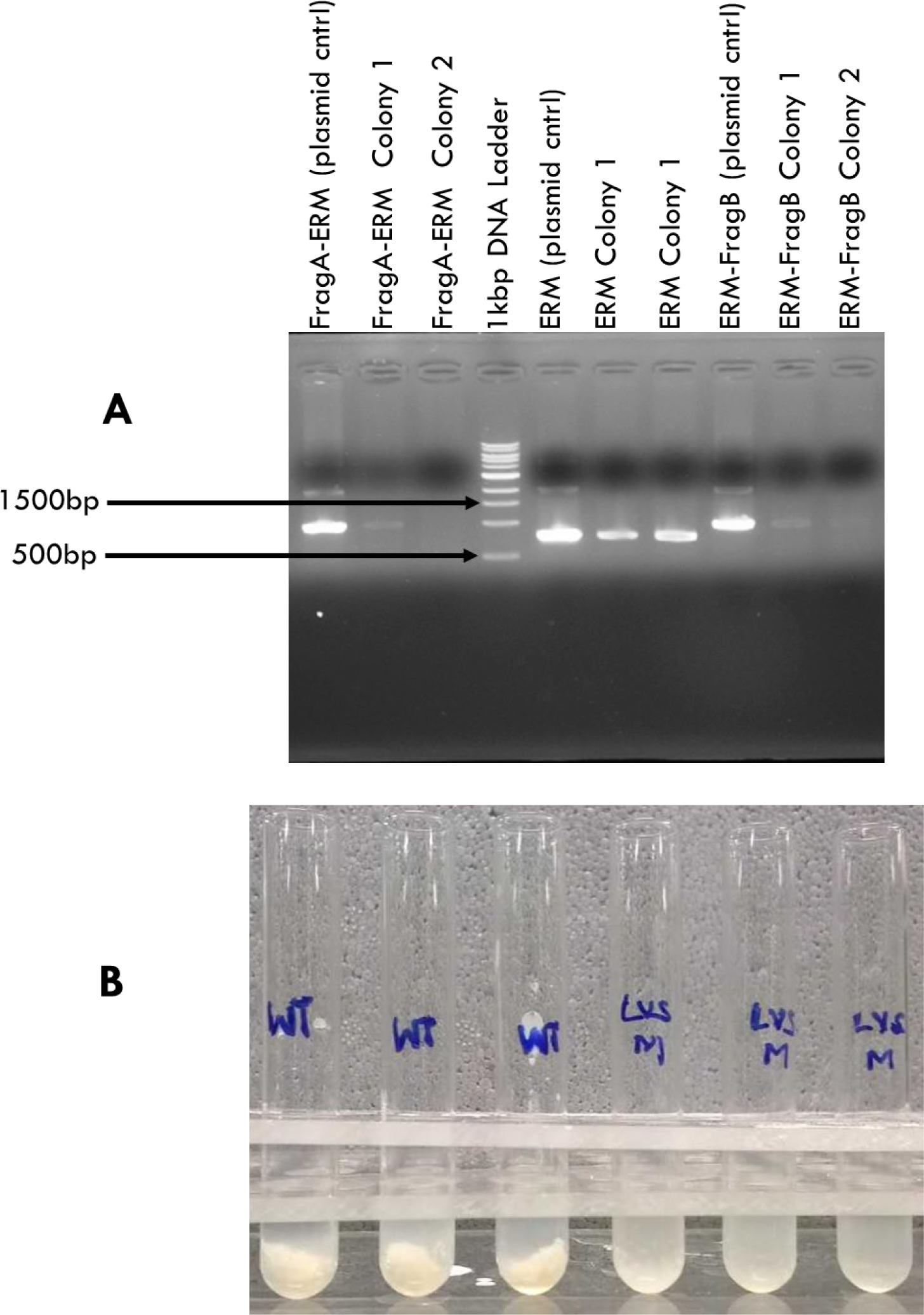
**A:** Gel image showing colony PCR of *P. polymyxa* levansucrase null mutant. Lanes show bands corresponding to LevFragA-ERM, ERM and ERM-LevfragB gene fragments. Other lanes show 1kb DNA ladder and pGEM7Zf+ harboring the levansucrase inactivation construct (bands from colonies 1 and 2). **B:** Precipitation of EPS in the fermentation broth of *P. polymyxa* wildtype and levansucrase null mutant in sucrose medium.

### Effect of levansucrase disruption on EPS formation

Batch 2,3-BD fermentations were conducted with sucrose and glucose substrates to evaluate EPS formation by the levansucrase null mutant. The fermentation cultures were supplemented with 0, 0.2, 0.4 g/L CaCl_2_. Fermentation on sucrose showed that EPS formation by the levansucrase null mutant decreased 5.8-, 6.4- and 6.1-fold in the 0, 0.2 and 0.4 g/L CaCl_2_-supplemented cultures, respectively, when compared to the wildtype (**Table 4, Fig. 2B**). The levansucrase null mutant showed no measurable levansucrase activity whereas more than 0.6 units of levansucrase activity per milligram protein were detected in the wildtype (**Figs 3F**, **4F**, **5F**). The absence of any measurable levansucrase activity in the mutant confirms successful inactivation of levansucrase gene in *P. polymyxa*. However, even though no measurable levansucrase activity was detected in the mutant, 2-3 g/L EPS was synthesized by the mutant grown on sucrose (**Table 4**), thus suggesting that *P. polymyxa* produces other EPS forms other than levan.

**Table 4.**
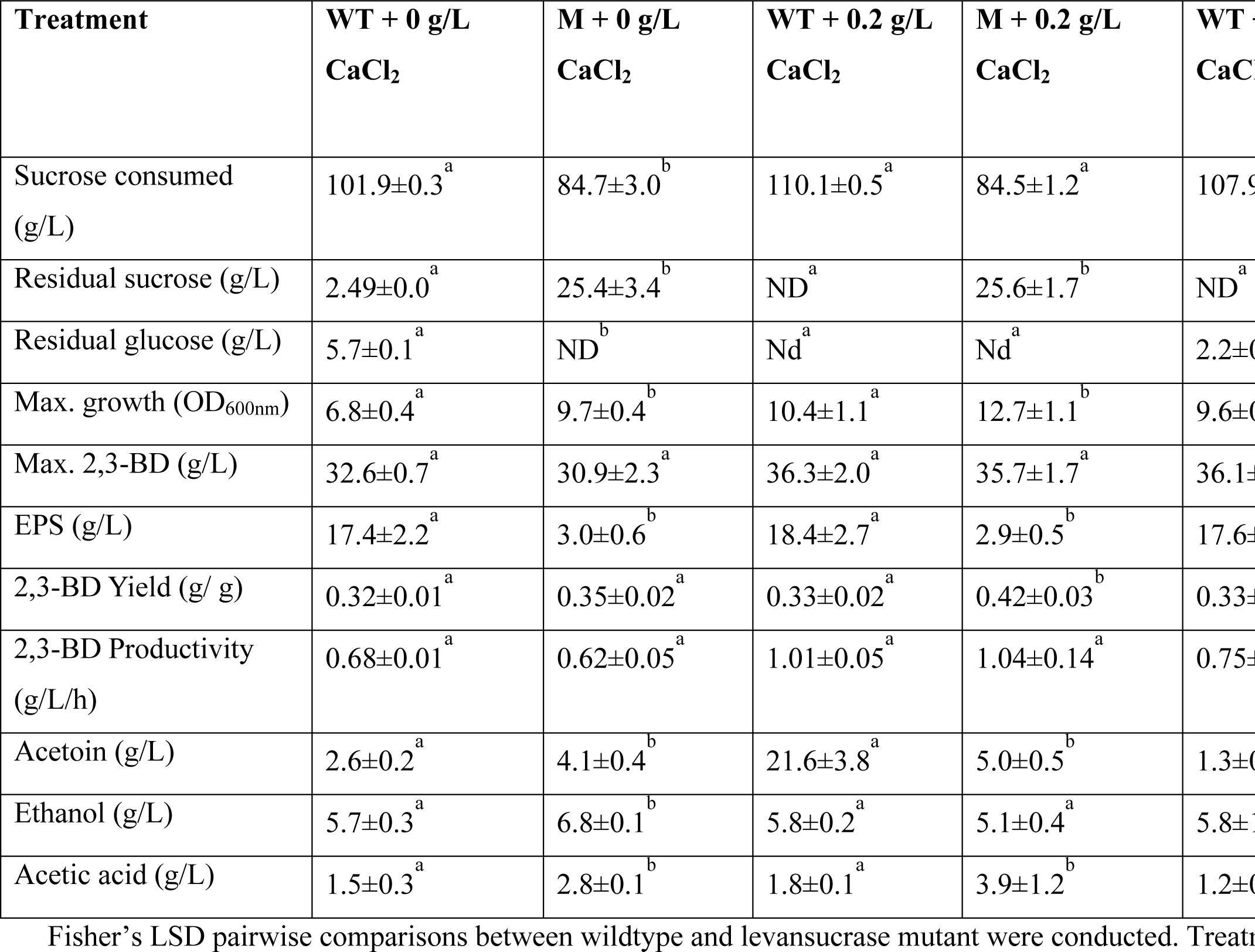
Substrate consumed, growth, maximum products, 2,3-BD yield and productivity during **sucrose** fermentation by *P. polymyxa* DSM 365 wildtype and levansucrase null mutant.

| Treatment | WT + 0 g/L<br>CaCl<sub>2</sub> | M + 0 g/L<br>CaCl<sub>2</sub> | WT + 0.2 g/L<br>CaCl<sub>2</sub> | M + 0.2 g/L<br>CaCl<sub>2</sub> | WT + 0.4 g/L<br>CaCl<sub>2</sub> | M + 0.4 g/L<br>CaCl<sub>2</sub> |
| --- | --- | --- | --- | --- | --- | --- |
| Sucrose consumed (g/L) | 101.9±0.3<sup>a</sup> | 84.7±3.0<sup>b</sup> | 110.1±0.5<sup>a</sup> | 84.5±1.2<sup>a</sup> | 107.9±0.4<sup>a</sup> | 92.8±1.3<sup>b</sup> |
| Residual sucrose (g/L) | 2.49±0.0<sup>a</sup> | 25.4±3.4<sup>b</sup> | ND<sup>a</sup> | 25.6±1.7<sup>b</sup> | ND<sup>a</sup> | 17.3±1.8<sup>b</sup> |
| Residual glucose (g/L) | 5.7±0.1<sup>a</sup> | ND<sup>b</sup> | Nd<sup>a</sup> | Nd<sup>a</sup> | 2.2±0.0<sup>a</sup> | ND<sup>b</sup> |
| Max. growth (OD<sub>600nm</sub>) | 6.8±0.4<sup>a</sup> | 9.7±0.4<sup>b</sup> | 10.4±1.1<sup>a</sup> | 12.7±1.1<sup>b</sup> | 9.6±0.9<sup>a</sup> | 12.8±0.9<sup>b</sup> |
| Max. 2,3-BD (g/L) | 32.6±0.7<sup>a</sup> | 30.9±2.3<sup>a</sup> | 36.3±2.0<sup>a</sup> | 35.7±1.7<sup>a</sup> | 36.1±2.5<sup>a</sup> | 37.4±0.9<sup>a</sup> |
| EPS (g/L) | 17.4±2.2<sup>a</sup> | 3.0±0.6<sup>b</sup> | 18.4±2.7<sup>a</sup> | 2.9±0.5<sup>b</sup> | 17.6±0.36<sup>a</sup> | 2.9±0.2<sup>b</sup> |
| 2,3-BD Yield (g/ g) | 0.32±0.01<sup>a</sup> | 0.35±0.02<sup>a</sup> | 0.33±0.02<sup>a</sup> | 0.42±0.03<sup>b</sup> | 0.33±0.02<sup>a</sup> | 0.40±0.02<sup>a</sup> |
| 2,3-BD Productivity (g/L/h) | 0.68±0.01<sup>a</sup> | 0.62±0.05<sup>a</sup> | 1.01±0.05<sup>a</sup> | 1.04±0.14<sup>a</sup> | 0.75±0.05<sup>a</sup> | 0.78±0.02<sup>a</sup> |
| Acetoin (g/L) | 2.6±0.2<sup>a</sup> | 4.1±0.4<sup>b</sup> | 21.6±3.8<sup>a</sup> | 5.0±0.5<sup>b</sup> | 1.3±0.4<sup>b</sup> | 6.0±0.6<sup>b</sup> |
| Ethanol (g/L) | 5.7±0.3<sup>a</sup> | 6.8±0.1<sup>b</sup> | 5.8±0.2<sup>a</sup> | 5.1±0.4<sup>a</sup> | 5.8±1.1<sup>a</sup> | 5.5±0.5<sup>a</sup> |
| Acetic acid (g/L) | 1.5±0.3<sup>a</sup> | 2.8±0.1<sup>b</sup> | 1.8±0.1<sup>a</sup> | 3.9±1.2<sup>b</sup> | 1.2±0.3<sup>a</sup> | 5.5±0.2<sup>b</sup> |

**Figure 3:**
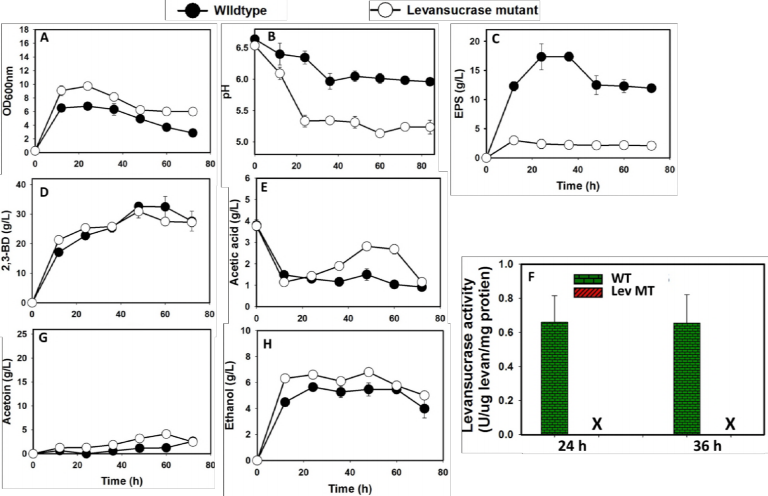
Fermentation profile of *P. polymyxa* levansucrase null mutant and wildtype grown in **sucrose** medium without CaCl_2_ supplementation. X represents no levansucrase activity. *NB: the underlined sequences represent either restriction sites or ribosomal binding site*

**Figure 4:**
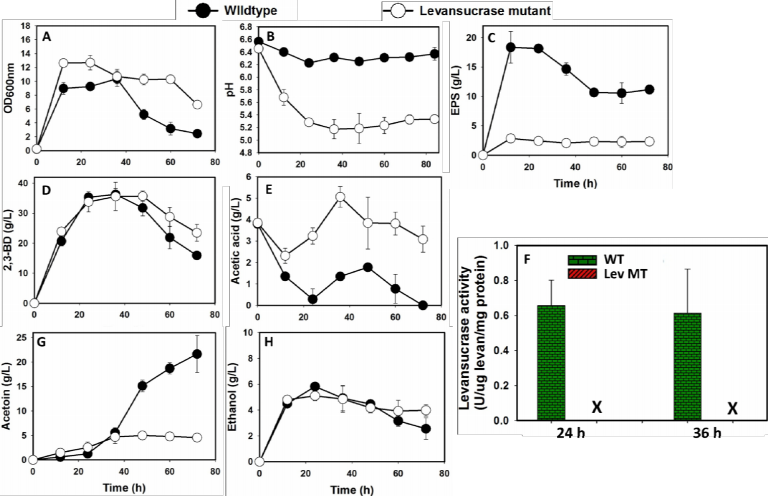
Fermentation profile of *P. polymyxa* levansucrase null mutant and wildtype grown in **sucrose** medium supplemented with 0.2 g/L CaCl_2._ X represents no levansucrase activity.

**Figure 5:**
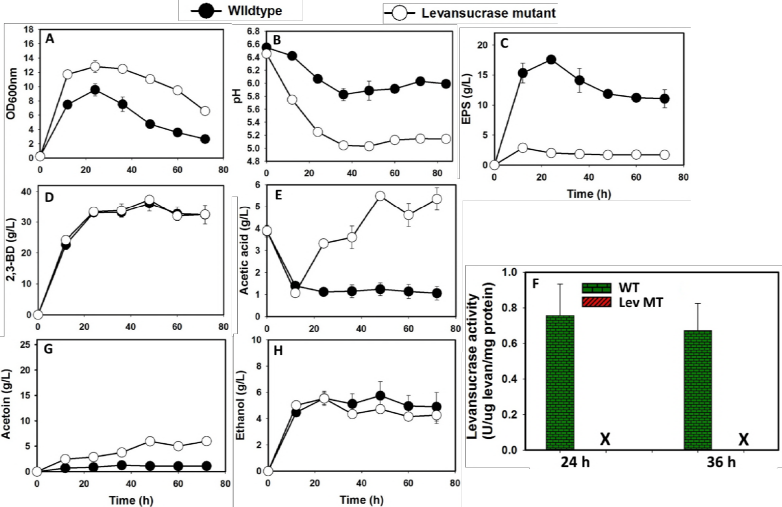
Fermentation profile of *P. polymyxa* levansucrase null mutant and wildtype grown in **sucrose** medium supplemented with 0.4 g/L CaCl_2._ X represents no levansucrase activity.

EPS production by the levansucrase null mutant grown on glucose decreased 2.4-, 1.7- and 1.9-fold in the 0, 0.2 and 0.4 g/L CaCl_2_-supplemented cultures, respectively, when compared to the wildtype (**Table 5; Figs S3-S5**). Interestingly, no levansucrase activity was observed in both the wildtype and the levansucrase null mutant grown on glucose. However, EPS produced by the wildtype cultures in the glucose medium decreased by at least 4-fold when compared to the cultures grown on sucrose (**Tables 4** and **5**).

**Table 5.** Substrate consumed, growth, maximum products, 2,3-BD yield and productivity during **glucose** fermentation by *P. polymyxa* DSM 365 wildtype and levansucrase null mutant.

| Treatment | WT + 0 g/L<br>CaCl<sub>2</sub> | M + 0 g/L<br>CaCl<sub>2</sub> | WT + 0.2 g/L<br>CaCl<sub>2</sub> | M + 0.2 g/L<br>CaCl<sub>2</sub> | WT + 0.4 g/L<br>CaCl<sub>2</sub> | M + 0.4 g/L<br>CaCl<sub>2</sub> |
| --- | --- | --- | --- | --- | --- | --- |
| Glucose consumed (g/L) | 85.6±3.9<sup>a</sup> | 87.8±1.9<sup>a</sup> | 96.1±3.1<sup>a</sup> | 105.0±0.0<sup>a</sup> | 94.7±2.7<sup>a</sup> | 106.7±2.5<sup>b</sup> |
| Residual glucose (g/L) | 23.7±3.2<sup>a</sup> | 21.45±1.2<sup>a</sup> | 13.2±2.3<sup>a</sup> | 4.3±0.7<sup>b</sup> | 14.5±2.0<sup>a</sup> | 2.5±1.8<sup>b</sup> |
| Max. growth (OD<sub>600nm</sub>) | 8.9±0.2<sup>a</sup> | 12.1±1.3<sup>b</sup> | 11.2±0.2<sup>a</sup> | 15.4±1.2<sup>b</sup> | 10.5±0.3<sup>a</sup> | 16.1±0.7<sup>b</sup> |
| Max. 2,3-BD (g/L) | 30.7±1.2<sup>a</sup> | 29.8±2.0<sup>a</sup> | 34.2±2.7<sup>a</sup> | 39.0±1.3<sup>a</sup> | 34.5±3.4<sup>a</sup> | 39.4±4.4<sup>a</sup> |
| EPS (g/L) | 5.5±0.4<sup>a</sup> | 2.27±0.1<sup>b</sup> | 4.2±0.2<sup>a</sup> | 2.4±0.3<sup>b</sup> | 5.4±0.5<sup>a</sup> | 2.8±0.2<sup>b</sup> |
| 2,3-BD Yield (g/ g) | 0.36±0.01<sup>a</sup> | 0.34±0.02<sup>a</sup> | 0.36±0.02<sup>a</sup> | 0.37±0.01<sup>a</sup> | 0.37±0.04<sup>a</sup> | 0.38±0.04<sup>a</sup> |
| 2,3-BD Productivity (g/L/h) | 0.51±0.02<sup>a</sup> | 0.57±0.09<sup>a</sup> | 0.71±0.06<sup>a</sup> | 1.62±0.05<sup>b</sup> | 0.72±0.07<sup>a</sup> | 1.64±0.19<sup>b</sup> |
| Acetoin (g/L) | 3.3±0.0<sup>a</sup> | 2.4±0.4<sup>b</sup> | 10.1±2.1<sup>a</sup> | 2.8±0.4<sup>b</sup> | 8.8±1.1<sup>a</sup> | 2.6±0.3<sup>b</sup> |
| Ethanol (g/L) | 5.8±0.1<sup>a</sup> | 7.1±0.3<sup>b</sup> | 5.6±0.4<sup>a</sup> | 8.7±0.7<sup>b</sup> | 5.4±0.5<sup>a</sup> | 9.4±0.2<sup>b</sup> |
| Acetic acid (g/L) | 0.0±0.0<sup>a</sup> | 1.7±0.0<sup>b</sup> | 0.6±0.0<sup>a</sup> | 2.5±0.9<sup>b</sup> | 0.0±0.0<sup>b</sup> | 2.9±0.1<sup>b</sup> |

### Effect of calcium supplementation on growth, sugar utilization, 2,3-BD yield and productivity

Initial fermentations with the levansucrase null mutant resulted in a sharp drop in pH, which adversely affected cell growth and product formation, particularly when the pH fell below 5.5 (**Figs 3A, B** and **S3A**, **B**). Thus, medium supplementation with CaCO_3_ and CaCl_2_ was adopted given pre-reported capacity of calcium to influence key cellular processes such as sugar transport, product formation and tolerance, and most importantly pH stabilization (Han et al., 2013; Okonkwo et al., 2016; Zeng et al., 2010). Medium supplementation with CaCO_3_ and CaCl_2_ did not improve culture pH of the levansucrase null mutant (data not shown), however, CaCl_2_ exerted remarkable influence on cell growth and 2,3-BD production by this strain. Following CaCl_2_ supplementation, cell biomass production increased in the sucrose- and glucose-grown cultures for both the wildtype and the levansucrase null mutant with attendant increases in substrate consumption (**Tables 4** and **5**). Growth of the levansucrase null mutant on sucrose increased by 22% and 34%, respectively, with0.2and 0.4 g/L CaCl_2_ supplementation when compared to the wildtype (**Figs 4A and 5A**). As shown in **Figs 4A** and **5A**, CaCl_2_ supplementation led to increased biomass accumulation. With CaCl_2_ (0.2 and 0.4 g/L), 2,3-BD yield on sucrose increased 27% in the levansucrase null mutant relative to the wildtype (**Table 4**). In addition, the 2,3-BD productivity of the null mutant on sucrose increased marginally - approximately 3% and 4% with 0.2 and 0.4 g/L CaCl_2_ treatments, respectively, when compared to the wildtype (**Table 4**). Conversely, without CaCl_2_ supplementation productivity of the levansucrase null mutant on sucrose decreased 8.8% compared to the wildtype (**Table 4**).

With glucose, addition of 0.2 and 0.4 g/L CaCl_2_ to cultures of the levansucrase null mutant increased growth by 27% and 34%, respectively, compared to the null mutant grown in cultures without CaCl_2_ supplementation (**Table 5**). Similarly, the wildtype exhibited 25% and 17% increased growth with 0.2 and 0.4 g/L CaCl_2_ relative to CaCl_2_-unsupplemented cultures (**Table 5**). As observed in the sucrose cultures, glucose utilization by the wildtype and levansucrase null mutant improved with 0.2 and 0.4 g/L CaCl_2_ supplementation. Glucose utilization by the mutant increased 20% and 22% in the 0.2 and 0.4 g/L CaCl_2_-supplemented cultures, respectively, when compared to the null mutant cultures without CaCl_2_ supplementation (**Table 5**). The 2,3-BD yield and productivity of the levansucrase null mutant grown on glucose increased from 0.34 g/g and 0.57 g/L/h (without CaCl_2_), respectively, to at least 0.37 g/g and 1.62 g/L/h (with CaCl_2_ supplementation), respectively, whereas, 2,3-BD yield and productivity of the wildtype increased from 0.36 g/g and 0.51 g/L/h (without CaCl_2_), respectively, to at least 0.36 g/g and 0.71 g/L/h (with CaCl_2_ supplementation), respectively. However, the 2,3-BD yield and productivity of the levansucrase null mutant in glucose cultures increased by 3% and 4% (without CaCl_2_), and 2% and 128%, respectively, in the CaCl_2-_supplemented cultures, respectively, relative to the wildtype (**Table 5**). The observed increased glucose utilization rate of the levansucrase null mutant may be responsible for the enhanced 2,3-BD titer, yield and productivity relative to the wildtype.

The levansucrase null mutant efficiently converted sucrose to 2,3-BD with diminished ability to produce EPS (**Fig. 2B**). However, the mutant utilized glucose much faster than sucrose resulting in higher 2,3-BD productivity on glucose relative to fermentations conducted with sucrose as substrate (**Tables 4** and **5**). The null mutant achieved a maximum 2,3-BD yield of 0.42 g/g with CaCl_2_ supplemented (0.2 g/L) when grown on sucrose, which is 27% more than that (0.33 g/g) achieved by the wildtype (**Tables 4**).

### qRT-PCR

To further confirm successful inactivation of levansucrase in *P. polymyxa*, qRT-PCR was conducted to quantify the mRNA transcript levels of levansucrase and erythromycin resistance genes of levansucrase null mutant relative to the wildtype. For these analyses, both strains were grown on sucrose and glucose. As shown in **Fig. 6**, mRNA transcripts of the *P. polymyxa* levansucrase gene were not detected in the null mutant grown on both sucrose and glucose. Conversely, we detected high levels of mRNA transcripts of the erythromycin resistance gene in the levansucrase null mutant (**Fig. 6**). Clearly, this indicates effective levansucrase inactivation and successful integration of the deletion construct in the genome of *P. polymyxa* by double cross homologous recombination.

**Figure 6.**
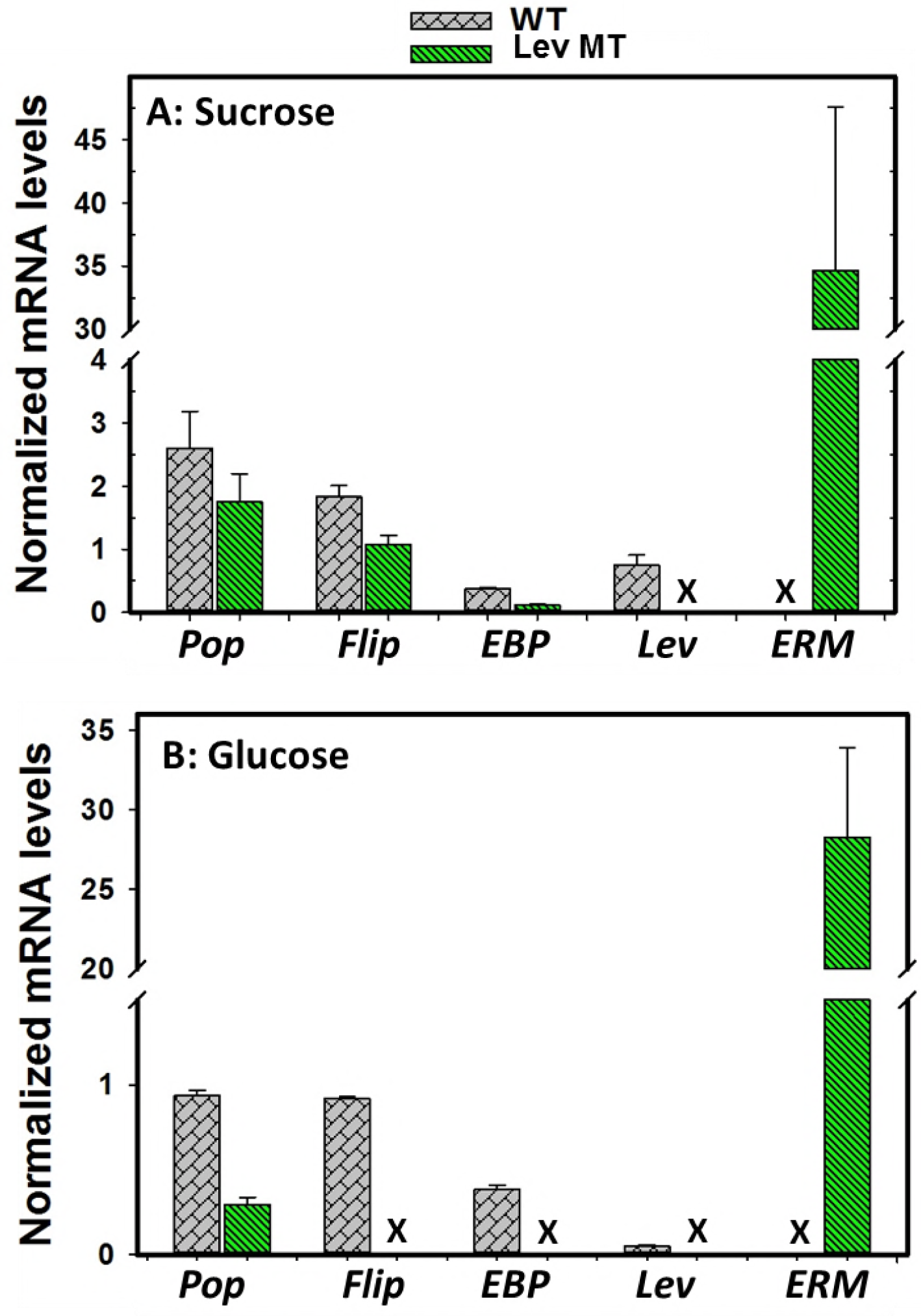
Comparisons of mRNA transcript levels of polysaccharide polymerase (*Pop*), flippase (*flip*), exopolysaccharide biosynthesis protein (*EBP*), levansucrase (*lev*), and erythromycin (*ERM*) genes of *P. polymyxa* wildtype (WT) and levansucrase null mutant (Lev MT) grown in sucrose (**A**) and glucose (**B**) media. X represents no mRNA detection.

Despite successful levansucrase gene inactivation in *P. polymyxa*, EPS accumulation was not completely abolished in cultures of the levansucrase null mutant. To delineate the source(s) of the observed EPS in the levansucrase null mutant, qRT-PCR was used to probe the mRNA transcript levels of other exopolysaccharide biosynthesis genes in *P. polymyxa*, namely, polysaccharide polymerase (*pop*), flippase (*flip*) and exopolysaccharide biosynthesis protein (*EBP*). *Pop, flip* and *EBP* genes are putative genes that have been implicated in the biosynthesis of other EPSs other than levan in *P. polymyxa* (35). *Pop, flip* and *EBP* genes were found to be expressed by the levansucrase null mutant grown in sucrose medium, however, only *Pop* was expressed by the levansucrase null mutant on glucose substrate (**Fig. 6**). *Pop* and *EBP* are putative genes that are possibly associated with exopolysaccharide precursors and assemble monomers to growing EPS chain (35). Different studies suggest that flippase might be membrane specific, where it assembles EPS repeating units and then translocates complete EPS across the membrane (35). The expression of *Pop, flip* and *EBP* suggests that *P. polymyxa* has the capability to synthesize other forms of EPS when grown on both sucrose and glucose.

### Stability of levansucrase null mutant

Stability of the levansucrase null mutant was tested over 50 generations. For this, 2,3-BD fermentations were conducted with different (0, 10, 20, 30, 40 and 50) generations of *P. polymyxa* levansucrase null mutant on sucrose substrate, with and without erythromycin supplementation. As shown in **Figs S6** and **S7**, the maximum growth for each tested generation of levansucrase null mutant under erythromycin pressure was between 11.0 and 13.9 (OD_600nm_), whereas without erythromycin, the growth ranged from 12.5 to 15.1 (OD_600nm_).

EPS concentrations at each tested generation (G_0_-G_50_) of the levansucrase null mutant, with and without erythromycin supplementation were considerably similar. The maximum 2,3-BD produced by the mutant grown under antibiotic pressure was in the range of 35.3 to 39.4 g/L, whereas without antibiotics, the 2,3-BD production was in the range of 36.1 to 39.0 g/L (**Table S1**). Colony-PCR and replica plating techniques were further employed to characterize the levansucrase null mutant generation (G_0_-G_50_) for antibiotic resistances. The results are shown in **Figs S8** and **S9**. Notably, the levansucrase null mutant developed in this study retained antibiotic resistance to erythromycin after 50 generations of growth with or without erythromycin addition.

## Discussion

EPS production during 2,3-BD fermentation constitutes a nuisance during fermentation and diverts substrate carbons away from 2,3-BD biosynthesis, thus decreasing 2,3-BD yield and productivity. Also, viscosity of the fermentation broth increases with EPS production, which impairs mixing (9). More importantly, EPS negatively impacts 2,3-BD downstream processing, thereby increasing the overall cost of production. Therefore, this study was aimed at developing a mutant strain of *P. polymyxa* with diminished ability to synthesize EPS. We employed double cross homologous recombination strategy to inactivate levansucrase gene in *P. polymyxa*. The following objectives were achieved: (i) disruption of levansucrase gene in *P. polymyxa* via erythromycin gene insertion, (ii) phenotypic characterization of the levansucrase null mutant by determining the resultant cell growth, and concentrations of EPS, 2,3-BD, acetoin, ethanol and acetic acid.

The genome of *P. polymyxa* DSM 365 is not fully sequenced. Thus, due to insufficient genomic information on this microorganism, the nucleotide and protein sequences of the *P. polymyxa* DSM 365 levansucrase from the available shot-gun sequences were compared to those of other *P. polymyxa* strains whose complete genome sequences are available on public databases. As shown in **Table 1**, fully sequenced *P. polymyxa* strains have a single copy of the levansucrase gene, which has 95-97% protein sequence similarity to that of *P. polymyxa* DSM 365. Levansucrase, which is responsible for levan EPS biosynthesis in *P. polymyxa* (36, 37), was targeted for inactivation. The results from the present study are discussed below.

### Effect of levansucrase inactivation on EPS biosynthesis by *P. polymyxa* levansucrase null mutant

Successful knockout of levansucrase gene in *P. polymyxa* was confirmed by PCR, restriction digest analysis, levansucrase activity assay, quantitative real-time PCR, antibiotic selection and genetic stability. Knockout of levansucrase gene in *P. polymyxa* resulted in significant reduction in EPS formation by the levansucrase null mutant grown on sucrose and glucose (**Figs. 3C, 4C, 5C, S3 C, S4 C, S5 C**). Clearly, significant reduction in the amount of EPS accumulated by the levansucrase null mutant confirms that levansucrase is the key player in EPS biosynthesis in *P. polymyxa* and that the targeted open reading frame (ORF) in the *P. polymyxa* DSM 365 shotgun sequence encodes levansucrase. Reduction in EPS biosynthesis by the levansucrase null mutant was more pronounced during growth on sucrose relative to glucose. This is ascribable to the fact that EPS formation is more strongly favored by sucrose, which is hydrolyzed by levansucrase to release glucose and fructose (38). The vast majority of the fructose molecules are then linked to form EPS by the same enzyme (levansucrase). Therefore, sucrose consumption by *P. polymyxa* results in significantly higher EPS production, which was almost completely abolished in the null mutant. In fact, levansucrase mRNA and activity were not detected in the levansucrase null mutant grown on both glucose and sucrose. However, levansucrase mRNA expression and activity were observed in the wildtype grown in sucrose-based medium. Levansucrase mRNA transcripts, albeit marginal, were detected in the wildtype cultures grown on glucose without any measurable levansucrase activity (**Fig. 6**). The lack of levansucrase activity in the wildtype in glucose medium coupled with the detection of low levels of levansucrase mRNA transcripts in the corresponding cells suggests that sucrose likely plays a specific role in levansucrase gene expression. In fact, sucrose-mediated induction of levansucrase gene expression has been reported by previous authors (39, 40). However, levansucrase mRNA transcript levels were not quantified in these studies. Notably, (41) observed levansucrase mRNA transcripts in *Z. mobilis* cultures on both glucose and fructose. Perhaps, different mechanisms govern levansucrase expression in different species, which employ EPS for different functions in their different ecological habitats/niches. Further studies are required to better delineate the role of sucrose in levansucrase gene expression in *P. polymyxa*.

Despite levansucrase gene inactivation which was confirmed by PCR, antibiotic sensitivity assay and replica plating, and qPCR, EPS was detected in cultures of both the wildtype and the levansucrase null mutant grown on glucose, albeit in significantly low amounts in the levansucrase null mutant. We therefore rationalized that *P. polymyxa* likely produces more than one type of extracellular polysaccharide with different sugars, with sucrose favoring levan biosynthesis, while glucose supports the production of uncharacterized polysaccharide(s). This is not unusual among EPS-producing microorganisms as *Bacillus* spp., *Z. mobilis, Leuconostoc mesenteriodes, Agrobacterium radiobacter, Xanthamonas campestris,* and *Pseudomonas aeruginosa* have been shown to produce alginate, xanthan, curdlan or dextran with different sugars (42, 43, 44, 45, 46, 47, 48). To test the likelihood that *P. polymyxa* produces another EPS other than levan, we used qPCR to assay for mRNA transcripts of genes likely involved in the expression of other EPSs. Interestingly, polysaccharide polymerase, flippase, and exopolysaccharide biosynthesis protein, which are putatively involved in the synthesis of other EPS, were expressed in both the levansucrase null mutant and the wildtype (**Figs 6** and **7**). The expression of polysaccharide polymerase gene by *P. polymyxa* levansucrase null mutant cultures in both sucrose and glucose indicates that other genes may be involved in the biosynthesis of other forms of EPS by *P. polymyxa* (**Fig 7**). Hence, physicochemical characterization of the EPS obtained in glucose cultures of *P. polymyxa* may shed more light on the other EPS forms.

**Figure 7.**
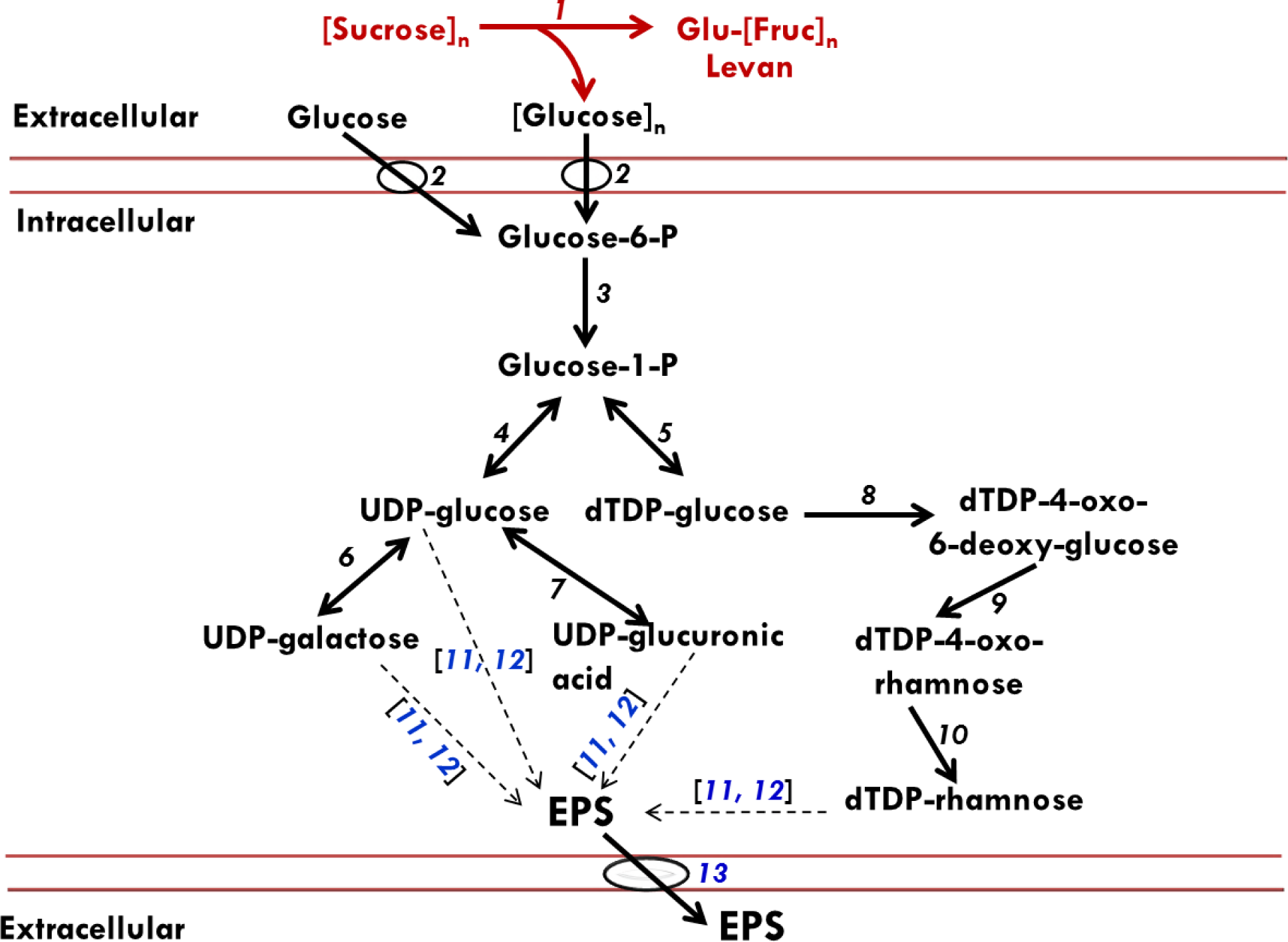
Schematic representation of annotated and putative pathways of EPS production by *P. polymyxa*. The schematic diagram was conceived based on our enzymatic assays, qRT-PCR data and annotated metabolic network model of *P. polymyxa* in KEGG databases. The red and black fonts or lines represent annotated and putative EPS production pathways in *P. polymyxa*, respectively. 1, levansucrase; 2, sugar PTS; 3, phosphoglucomutase; 4, UTP-glucose 1-phosphate uridylyltransferase; 5, glucose 1-phosphate thymidylyltransferase; 6, galactose -1-phosphate uridylyltransferase; 7, UDP-glucose 6-dehydrogenase; 8, dTDP-glucose 4,6-dehydratase; 9, dTDP-4-dehydrorhamnose 3,5-epimerase; 10, dTDP-4-dehydrorhamnose reductase; 11, polymerase synthase; 12, exopolysaccharide biosynthesis protein; 13, flippase. The broken black arrows represent EPS polymerization steps from sugar nucleotides. The genes that code the enzymes (numbers 11, 12, and 13) in blue font were analyzed by qRT-PCR.

Whereas EPS production was significantly reduced in the levansucrase mutant when compared to the wildtype, this did not translate to significant increase in 2,3-BD production with sucrose as substrate. In parallel, while sucrose utilization by the levansucrase null mutant was 1.3- fold lower than that of the wildtype, the amount of 2,3-BD produced was similar for both strains (**Table 4**). This implies that by utilizing considerably less substrate (sucrose), the levansucrase null mutant produced the same amount of product. This is an attractive trait (higher yield) for potential large-scale application from an economic standpoint. Further, reduced sucrose utilization by the levansucrase null mutant underscores disruption of sucrose utilization or processing following levansucrase inactivation in the null mutant. Comparatively, the levansucrase null mutant utilized 1.1-fold more glucose than the wildtype, which lends further weight to the role of levansucrase in sucrose utilization in *P. polymyxa* and successful inactivation of the encoding gene. Increased glucose utilization by the null mutant therefore accounts for 1.1-fold and 1.7-fold increases in 2,3-BD and ethanol production, when compared to the wildtype. Increased product accumulation (2,3-BD and ethanol) by the levansucrase mutant relative to the wildtype may stem from redirection of free carbons from EPS biosynthesis to the 2,3-BD and ethanol biosynthesis pathways. However, EPS accumulation was only slightly reduced in the levansucrase null mutant grown in glucose-based medium relative to the wildtype, so carbon redirection does not fully account for the increases in product formation. Therefore, it is likely that a different mechanism might be at play. Perhaps, levansucrase inactivation relieved a growth limiting machinery in the levansucrase null mutant leading to the observed increase in growth and consequently, increased product formation. Furthermore, ethanol production was clearly enhanced in the levansucrase null mutant relative to the wildtype when grown on glucose; an effect that was not observed with sucrose. It is not clear why this occurred, thus warranting further study. However, this result highlights ethanol biosynthesis as a veritable candidate for future inactivation towards developing a 2,3-BD over-producing strain.

### Levansucrase inactivation in *P. polymyxa* interferes with acetic acid re-assimilation

The *P. polymyxa* levansucrase null mutant was characterized for growth by measuring optical density (OD_600nm_) during 2,3-BD fermentation. Our results clearly suggest that levansucrase inactivation did not significantly impair the growth of *P. polymyxa* (**Tables 4** and **5**). Initial experiments showed low culture pH stemming from the accumulation of acetic acid in cultures of the levansucrase null mutant (**Fig. 3E**). To mitigate pH-related stresses and increase growth and product formation, CaCl_2_ was added to *P. polymyxa* levansucrase null mutant and wildtype cultures. CaCl_2_ supplementation enhanced the growth, 2,3-BD production, sugar utilization, and tolerance to pH stresses in *P. polymyxa*, particularly, the levansucrase null mutant (**Figs. 3-5; S3-S5; Tables 4** and **5**).

Typically, 2,3-BD is produced via a mixed acid pathway, which results in the accumulation of acetic, formic and lactic acids during fermentation. However, acetic acid is re-assimilated during fermentation with concomitant increase in culture pH. We observed that inactivation of the levansucrase gene in *P. polymyxa* resulted in acetic acid accumulation, leading to a decrease in culture pH (**Figs 3B** and **S3 B**). The acetic acid profile of the levansucrase null mutant relative to the wildtype suggests that levansucrase may directly or indirectly influence acetic acid re-assimilation. In both the glucose and sucrose-based cultures, with or without CaCl_2_ supplementation, elevated acetic acid accumulation was observed for the null mutant (**Figs. 3E, 4E, 5E, S3 E, S4 E, and S5 E**). There are no previous reports on possible links between EPS biosynthesis and acetic acid assimilation in *P. polymyxa* or other 2,3-BD producers, hence, this finding warrants further examination. More importantly, this indicates that EPS biosynthesis might play broader roles in the biology of *P. polymyxa*, and perhaps other 2,3-BD producers, as most are known to accumulate EPS during fermentation. Further, it is likely that acetic acid accumulation in the levansucrase null mutant is a secondary or cascade effect stemming from downstream effectors of levansucrase not directly involved in EPS biosynthesis. Solvent-producing, biphasic Gram-positive bacteria typically produce acids and then, reabsorb them during solvent formation. Disruption of their native biology has been reported to engender acid accumulation, due to poor acid assimilation. A similar pattern has been previously reported for *C. beijerinckii* NCIMB 8052 following knockdown of acetoacetate decarboxylase (49) and *Clostridium acetobutylicum* ATCC 842 (50). Perhaps, a similar phenomenon exists in *P. polymyxa* (which is also a solvent-producing biphasic Gram-positive bacterium). Notably, despite acetic acid accumulation and the attendant drop in culture pH, the levansucrase null mutant exhibited an overall improved growth than the wildtype in all conditions tested. It therefore appears that levansucrase inactivation might confer some form of stress resistance on the mutant cells which mitigates acid-mediated stress.

Overall, CaCl_2_ supplementation enhanced the growth of the mutant and the wildtype on both glucose and sucrose. This is attributable to previously reported global effects of calcium on cellular metabolism, sugar utilization and stress mitigation including upregulation of heat shock proteins (involved in the repair of damaged or aberrant proteins) and DnaK involved in DNA synthesis, transcription and repair in *C. beijerinckii* NCIMB 8052 (51). Additionally, Ca^2+^ has been implicated in the stabilization of bacterial membrane, which reduces the effects of membrane-damaging factors such as acids (52, 53). Therefore, the effects observed with CaCl_2_ for both strains, albeit more pronounced in the levansucrase null mutant of *P. polymyxa*, in which acetic acid accumulation was evident, likely stemmed from Ca^2+^-mediated mitigation of pH stresses.

### Stability of *P. polymyxa* levansucrase null mutant

The stability of microbial strains intended for industrial bioprocesses is critical for uniform and consistent product generation. This is particularly important when genetically modified strains are used. Hence, stability of the levansucrase null mutant generated in this study was tested. Stability results clearly showed that this mutant is stable as demonstrated by similar fermentation profiles (growth, 2,3-BD concentration, acid and ethanol concentrations, and EPS production) for the levansucrase null mutant grown to different generation times (up to 50 generations) in the presence and absence of antibiotic (**Figs S6** and **S7**).

## Conclusions

Levansucrase gene was successfully inactivated in *P. polymyxa* via double cross homologous recombination, leading to a stable and faster-growing strain with significantly reduced EPS production, which constitutes a significant nuisance to 2,3-BD fermentation and product recovery. The ability of the levansucrase null mutant to produce higher concentrations of 2,3-BD on glucose makes it attractive as a basis for generating a 2,3-BD overproducing strain from lignocellulosic biomass; of which glucose is the major sugar component. Towards further increasing 2,3-BD titer, yield and productivity, inactivation of ethanol biosynthesis appears a rational target considering increased ethanol accumulation by the mutant, especially when grown on glucose. Culture supplementation with small amounts of CaCl_2_ has promise as a means of mitigating metabolic disruptions that might arise following metabolic engineering of *P. polymyxa*; a typical occurrence in solvent-producing Gram-positive bacteria.

## Acknowledgements

Salaries and research support were provided in part by State funds appropriated to the Ohio Agricultural Research and Development Center (OARDC), OARDC graduate and interdisciplinary grants, and Hatch grant (Project No. OHO01222). We would like to also acknowledge Peloton Technologies LLC for financial support.

## References

1. Celinska E, Grajek W. 2009. Biotechnology production of 2, 3-butanediol-Current state and prospects. Biotechnol Adv 27: 715–725.

2. Ji X, Huang H, Ouyang P. 2011. Microbial 2, 3 – butanediol production: a state-of-the-art review. Biotechnol Adv 29: 351–364.

3. Syu MJ. 2001. Biological production of 2,3-butanediol. Appl Microbiol Biotechnol 55:10–8.

4. Garg S, Jain A. 1995. Fermentative production of 2,3-butanediol: a review. Bioresour Technol 51:103–109.

5. Guo X, Cao C, Wang Y, Li C, Wu M, Chen Y, Zhang C, Pei H, Xiao D. 2014. Effect of the inactivation of lactate dehydrogenase, ethanol dehydrogenase, and phosphotransacetylase on 2,3-butanediol production in Klebsiella pneumonia strain. Biotechnol Biofuels 7:44.

6. Zeng AP, Deckwer WD. 1991. A model for multiproduct-inhibited growth of Enterobacter aerogenes in 2, 3-butanediol fermentation. Appl Microbiol Biotechnol 35(1):1–3.

7. Okonkwo CC, Ujor V, Mishra PK, Ezeji TC. 2017. Process development for enhanced 2,3-butanediol production by Paenibacillus polymyxa DSM 365. Fermentation 3(2):18.

8. Priya A, Dureja P, Talukdar P, Rathi R, Lal B, Sarma PM. 2016. Microbial production of 2, 3-butanediol through a two-stage pH and agitation strategy in 150l bioreactor. Biochem Eng J 105:159–67.

9. Häßler T, Schieder D, Pfaller R, Faulstich M, Sieber V. 2012. Enhanced fed-batch fermentation of 2,3-butanediol by Paenibacillus polymyxa DSM 365. Bioresour Technol 124:237–244.

10. Ji XJ, Huang H, Du J, Zhu JG, Ren LJ, Li S, Nie ZK. 2009. Development of an industrial medium for economical 2, 3-butanediol production through co-fermentation of glucose and xylose by Klebsiella oxytoca. Bioresour Technol 100(21):5214–8.

11. Biebl H, Zeng AP, Menzel K, Deckwer WD. 1998. Fermentation of glycerol to 1,3-propanediol and 2,3-butanediol by Klebsiella pneumoniae. Appl Microbiol Biotechnol 50:24–9.

12. Jung MY, Ng CY, Song H, Lee J, Oh MK. 2012. Deletion of lactate dehydrogenase in Enterobacter aerogenes to enhance 2, 3-butanediol production. Appl Microbiol Biotechnol 95(2):461–9.

13. Jung MY, Mazumdar S, Shin SH, Yang KS, Lee J, Oh MK. 2014. Improvement of 2, 3-butanediol yield in Klebsiella pneumoniae by deletion of the pyruvate formate-lyase gene. Appl Environ Microbiol 80(19):6195–203.

14. Jantama K, Polyiam P, Khunnonkwao P, Chan S, Sangproo M, Khor K, Jantama SS, Kanchanatawee S. 2015. Efficient reduction of the formation of by-products and improvement of production yield of 2, 3-butanediol by a combined deletion of alcohol dehydrogenase, acetate kinase-phosphotransacetylase, and lactate dehydrogenase genes in metabolically engineered Klebsiella oxytoca in mineral salts medium. Metab Eng 30:16–26.

15. Nakashimada Y, Marwoto B, Kashiwamura T, Kakizono T, Nishio N. 2000. Enhanced butanediol production by addition of acetic acid in Paenibacillus polymyxa. J Biosci Bioeng 90:661–664.

16. De Mas CD, Jansen NB, Tsao GT. 1988. Production of optically active 2,3-butanediol by Bacillus polymyxa. Biotechnol Bioeng 31:366–77.

17. Donot F, Fontana A, Baccou JC, Schorr-Galindo S. 2012. Microbial exopolysaccharides: Main examples of synthesis, excretion, genetics and extraction. Carbohydr Polym 87: 951–962.

18. Liang TW, Wang SL. 2015. Recent advances in exopolysaccharides from Paenibacillus spp. production, isolation, structure, and bioactivities. Mar Drugs 13(4):1847–63.

19. Lal S, Tabacchioni S. 2009. Ecology and biotechnological potential of Paenibacillus polymyxa: a minireview. Indian J Microbiol 49(1):2–10.

20. Haggag WM. 2007. Colonization of exopolysaccharide-producing Paenibacillus polymyxa on peanut roots for enhancing resistance against crown rot disease. Afr J Biotechnol 6(13).

21. Lebuhn M, Heulin T, Hartmann A. 1997. Production of auxin and other indolic and phenolic compounds by Paenibacillus polymyxa strains isolated from different proximity to plant roots. FEMS Microbiol Ecol 22(4):325–34.

22. Sambrook J, Russel DW. 2001. Molecular Cloning, third ed. Cold Spring Harbor Laboratory Press, New York.

23. Yanisch-Perron C, Vieira J, Messing J. 1985. Improved M13 phage cloning vectors and host strains: nucleotide sequences of the M13amp18 and pUC19 vectors. Gene 33: 103–119.

24. Görke B, Stülke J. 2008. Carbon catabolite repression in bacteria: many ways to make the most out of nutrients. Nat Rev Microbiol 6(8):613–24.

25. Inukai M, Isono F, Takatsuki A. 1993. Selective inhibition of the bacterial translocase reaction in peptidoglycan synthesis by mureidomycins. Antimicrob Agents Chemother 37(5):980–983.

26. Zhou Y, Johnson EA. 1993. Genetic transformation of Clostridium botulinum by electroporation. Biotechnol Lett 15, 121–126.

27. Schmittgen TD, Livak KJ. 2008. Analyzing real-time PCR data by the comparative CT method. Nat Protoc 3:1101–1110.

28. Okonkwo CC, Ujor V, Ezeji TC. 2017. Investigation of the relationship between 2,3-butanediol toxicity and production during growth of Paenibacillus polymyxa. N Biotechnol 34(1):23–31.

29. Okonkwo CC, Azam MM, Ezeji TC, Qureshi N. 2016. Enhancing ethanol production from cellulosic sugars using Scheffersomyces (Pichia) stipitis. Bioprocess Biosyst Eng 39(7):1023–1032.

30. Zhang J, Wang R, Jiang P, Liu Z. 2002. Production of an exopolysaccharide bioflocculant by Sorangium cellulosum. Lett Appl Microbiol 34:178–81.

31. Nielsen SS. 2010. Phenol-sulfuric acid method for total carbohydrates. In Food Analysis Laboratory Manual, Springer USA. Pp. 47-53.

32. Dubois M, Gilles JK, Hamilton PA, Rebers PA, Smith F. 1956. Colorimetric method for determination of sugars and related substances. Anal Chem 28(3), 350–356.

33. Euzenat O, Guibert A, Combes D. 1997. Production of fructo-oligosaccharides by levansucrase from Bacillus subtilis C4. Process Biochem 2(3):237–243.

34. Bradford MM. 1976. A rapid and sensitive method for the quantitation of microgram quantities of protein utilizing the principle of protein-dye binding. Anal Biochem 72: 248–254.

35. Rutering M, Cress BF, Schilling M, Ruhmann B, Koffas MAG, Sieber V, Schmid J. 2017. Tailor-made exopolysaccharides-CRISPR-Cas9 mediated genome editing in Paenibacillus polymyxa. Synth Biol 2(1): ysx007.

36. Rütering M, Schmid J, Rühmann B, Schilling M, Sieber V. 2016. Controlled production of polysaccharides–exploiting nutrient supply for levan and heteropolysaccharide formation in Paenibacillus sp. Carbohydr Polym 148:326–34.

37. Choi HJ, Kim CS, Kim P, Jung HC, Oh DK. 2004. Lactosucrose bioconversion from lactose and sucrose by whole cells of Paenibacillus polymyxa harboring levansucrase activity. Biotechnol Prog 20(6):1876–9.

38. Yanase H, Iwata M, Nakahigashi R, Kita K, Kato N, Tonomura K. 1992. Purification, crystallization, and properties of the extracellular levansucrase from Zymomonas mobilis. Biosci Biotechnol Biochem 56(8):1335–7.

39. Arietta JG, Sotolongo M, Menendez C, Alfonso D, Trujillo LE, Soto M, Ramirez R, Hernandez L. 2004. A type II protein secretory pathway required for levansucrase secretion by Gluconacetobacter diazotrophicus. J Bacteriol 186:15031–5039.

40. Bezzate S, Aymerich S, Chambert R, Czarnes S, Berge O, Heulin T. 2000. Disruption of the Paenibacilus polymyxa levansucrase gene impairs its ability to aggregate soil in the wheat rhizosphere. Environ Microbiol 2:333–342.

41. Gurunathan S, Gunasekaran, P. 2004. Differential expression of Zymomonas mobilis sucrose genes (SacB and SacC) in Escherichia coli and sucrase mutants of Zymomonas mobilis. Braz Arch Biol Technol 47(3):329–338.

42. Koepsell HJ, Tsuchiya HM, Hellman NN, Kazenko, A, Hoffman CA, Sharpe ES, Jackson RW. 1953. Enzymatic synthesis of dextran acceptor specificity and chain initiation. J Biol Chem 200(2):793–801.

43. Esser K, Kadereit JW, Lüttge U, Runge M. 2012. Progress in Botany: Genetics Cell Biology and Physiology Systematics and Comparative Morphology Ecology and Vegetation Science. Springer Science & Business Media

44. Colvin KM, Irie Y, Tart CS, Urbano R, Whitney JC, Ryder C, Howell PL, Wozniak DJ, Parsek MR. 2012. The Pel and Psl polysaccharides provide Pseudomonas aeruginosa structural redundancy within the biofilm matrix. Environ Microbiol 14(8):1913–28.

45. Cooley BJ, Dellos-Nolan S, Dhamani N, Todd R, Waller W, Wozniak D, Gordon VD. 2016. Asymmetry and inequity in the inheritance of a bacterial adhesive. New J Phys 18(4):045019.

46. Papagianni M, Psomas SK, Batsilas L, Paras SV, Kyriakidis DA, Liakopoulou-Kyriakides M. 2001. Xanthan production by Xanthomonas campestris in batch cultures. Process Biochem 37(1):73–80.

47. Saudagar PS, Singhal RS. 2004 Fermentative production of curdlan. Appl Biochem Biotechnol 118(1):21–31.

48. Zhang Z, Chen H. 2010. Fermentation performance and structure characteristics of xanthan produced by Xanthomonas campestris with a glucose/xylose mixture. Appl Biochem Biotechnol 160(6):1653–63.

49. Han B, Gopalan V, Ezeji TC. 2011. Acetone production in solventogenic Clostridium species: new insights from non-enzymatic decarboxylation of acetoacetate. Appl Microbiol Biotechnol 91(3):565–76.

50. Ren C, Gu Y, Hu S, Wu Y, Wang P, Yang Y, Yang C, Yang, S, Jiang W. 2010. Identification and inactivation of pleiotropic regulator CcpA to eliminate glucose repression of xylose utilization in Clostridium acetobutylicum. Metab Eng 12(5):446–54.

51. Han B, Ujor V, Lai LB, Gopalan V, Ezeji TC. 2013. Use of proteomic analysis to elucidate the role of calcium in acetone-butanol-ethanol fermentation by Clostridium beijerinckii NCIMB 8052. Appl Environ Microbiol 79(1):282–93.

52. Hansen LT, Austin JW, Gill TA. 2001. Antibacterial effect of protamine in combination with EDTA and refrigeration. Int J Food Microbiol 66: 149 –161.

53. Kotra LP, Golemi D, Amro NA, Liu GY, Mobashery S. 1999. Dynamics of the lipopolysaccharides assembly on the surface of Escherichia coli. J Am Chem Soc 121:8707–8711.

